# Genetic Manipulation of Caveolin-1 in the Mouse Model of Marfan Syndrome Associated Aortic Root Aneurysm: Effects on Endothelial and Smooth Muscle Function

**DOI:** 10.1101/2024.07.29.605631

**Authors:** Tala Curry, Brikena Gusek, Ross M. Potter, T. Bucky Jones, Raechel Dickman, Nathan Johnson, Johana Vallejo-Elias, Mitra Esfandiarei

**Affiliations:** Biomedical Sciences Program, College of Graduate Studies, Midwestern University, Glendale, AZ; Department of Physiology, College of Graduate Studies, Midwestern University, Glendale, AZ; Department of Anatomy, College of Graduate Studies, Midwestern University, Glendale, AZ; Department of Anesthesiology, Pharmacology & Therapeutics, University of British Columbia, Vancouver, BC; Department of Basic Medical Sciences, College of Medicine, University of Arizona, Phoenix, AZ

**Keywords:** Marfan Syndrome, Aortic Aneurysm, Caveolin-1, Smooth Muscle, Endothelium

## Abstract

Marfan Syndrome (MFS) is a systemic connective tissue disorder caused by mutations in the gene encoding for the large glycoprotein Fibrillin-1 (Fbn1), leading to wide-spectrum clinical manifestations, with the most life-threatening being aortic root aneurysm. MFS aortic aneurysm is known to be associated with reduced endothelial nitric oxide synthase (eNOS)-mediated nitric oxide (NO) production. Previous studies have shown that caveolin-1 (Cav1), a coat protein of caveolae structure on the plasma membrane, acts as a negative regulator of eNOS activity. This suggests that Cav1 may play a role in the development of aortic root aneurysm in MFS by modulating eNOS activity. In this study, we investigated the role of Cav1 in regulating aortic smooth muscle and endothelial function, aortic wall elasticity, and wall strength by generating MFS mice (*FBN1^+/Cys1041Gly^*) lacking *Cav1* gene expression (MFS/*Cav1*KO). Our data show that ablation of the *Cav1* gene results in a significant decrease in aortic smooth muscle contraction in response to the vasoconstricting agent phenylephrine seemingly due to a marked increase in NO production within the aortic wall. We also showed that acetylcholine-induced vasorelaxation was increased in MFS/*Cav1*KO mice potentially through the endothelial nitric oxide-dependent mechanism, further confirming inhibitory role of Cav1 on endothelial NO production. In addition, aortic wall elastin fiber structure and strength were markedly improved in male MFS/*Cav1*KO mice. This study demonstrates the regulatory role of Cav1 during the development of aortic root aneurysm in MFS mice through its effects on smooth muscle and endothelial functions in an NO-dependent manner.

## INTRODUCTION

Marfan syndrome (MFS) is the most common monogenetic autosomal dominant disorder of connective tissue. MFS is caused by mutations in the gene encoding for the large extracellular matrix (ECM) glycoprotein fibrillin-1 (Fbn1). MFS has an estimated prevalence of 1 in 3000- 5000 with no biological sex or ethnic bias ^1,2^. Fbn1 is one of the major constituents of extracellular microfibrils found in connective tissues that provides scaffolding support for the formation and maturation of elastin fibers, thus, supporting the integrity and elasticity of connective tissues ^3,4^. In addition to its structural role, Fbn1 can regulate multiple signaling molecules within the extracellular matrix (ECM), particularly, the activity of the cytokine transforming growth factor beta (TGF-β) through its ability to sequester the latent form of TGF-β within the ECM ^5–7^. The sequestering function of Fbn1 is achieved by binding to latent TGF-β binding proteins (LTBPs) that are essential for the storage and/or timely release of latent TGF-β complexes ^5–7^. Mutations in the *Fbn1* gene can cause deficiencies in functional and structural capacities of microfibrils and subsequent increases in the bioavailable form of TGF-β, leading to over-activation of the TGF-β downstream signaling pathway ^8,9^.

Studies in human MFS patients and the experimental mouse model have shown that the progression of aortic root aneurysm is also associated with endothelial dysfunction, as evidenced by a significant decrease in endothelial nitric oxide synthase (eNOS) activity within the aortic wall ^10–12^. In our previous report, we were able to show that overexpression of a constitutively active form of eNOS could block the development of aortic root aneurysm in MFS mice ^13^.

In the context of vascular biology, eNOS activity is regulated by a complex network of multiple signaling pathways at both extracellular and intracellular levels, involving a long list of multiple stimuli and signaling cascades including shear stress ^14^, phosphoinositide 3-kinase/Akt ^15^, bradykinin ^16^, calcium-calmodulin ^17^, acetylcholine/G protein-coupled receptors ^18^, vascular endothelial growth factor (VEGF) ^19^, estrogen receptors ^20^, cyclic AMP ^21^, and caveolin-1 (Cav1)^22–24^.

Cav1 protein is the main structural component of caveolae on the plasma membrane of all cell types. It is particularly expressed on endothelial cells, smooth muscle cells, skeletal muscle cells, and adipocytes ^25,26^. Caveolae are invaginations of lipid rafts that are considered as highly dense regions enriched in proteins, cholesterol, and sphingolipids ^27^. The shape and density of these structures allows for many interactions between various signaling molecules and membrane receptors ^28,29^. As the main protein in caveolae, Cav1 plays a key role in multiple cellular processes and signaling involved in inflammation, cell migration, cell differentiation, vesicular trafficking, lipid homeostasis, and stress responses. In the vascular wall, Cav1 plays multiple regulatory roles that include controlling eNOS activity in the endothelial cells ^24,30,31^, regulating vascular smooth muscle cells proliferation and migration ^32,33^, facilitating cholesterol transport and homeostasis in adipose tissue ^34,35^, and modulating the activity of matrix metalloproteinases (MMPs) involved in wall remodeling ^36,37^.

In the endothelium, Cav1 negatively regulates eNOS through direct binding. Through the inhibitory effects on eNOS function, Cav1 plays a critical role in regulating vascular tone and blood pressure ^38^. Previous studies have shown that disruption of Cav1 expression results in dysregulated NO production, leading to hypertension and atherosclerosis ^7,39–41^.

It is well understood that MFS aortic root aneurysm is associated with endothelial dysfunction and a significant decrease in eNOS phosphorylation and activity ^11,13^. Considering the important role of Cav1 in negatively regulating eNOS function, we designed a study to investigate the effects of Cav1 on aortic wall endothelium and smooth muscle cells (SMCs) in a mouse model of MFS-associated aortic root aneurysm (*FBN1^+/Cys1041Gly^*) by generating male and female MFS mice lacking the *Cav1* gene (MFS/*Cav1*KO). In this study we have measured the impact of *Cav1* deletion on endothelium-dependent vasorelaxation, SMCs contraction, and aortic wall strength using the small vessel chamber myography technique. We also assessed aortic wall elastin structural integrity in cross sections of the ascending aorta of MFS mice using *van Gieson* elastin staining.

## MATERIALS & METHODS

### Experimental mouse model

The MFS mouse model is heterozygous for an *Fbn1* allele (*Fbn1^C1041G/+^*), replicating the phenotype of aortic root aneurysm associated with the most common mutation seen in MFS aneurysm patients.^21^ A breeding colony of MFS mice was established in the Midwestern University Animal Facility. The female *Cav1* KO mouse (B6.Cg-*Cav1^tm1Mls^*/J) was purchased (Jackson Laboratory, USA) and crossed with male MFS mouse (*Fbn1^C1041G/+^*) through multiple backcrossing to generate MFS mice lacking the *Cav1* gene (*FBN1^-/+^/Cav1^-/-^*) that are referred to MFS/*Cav1KO* throughout the report. The breeding protocol produced equal numbers of MFS/*Cav1KO* and *Cav1* KO mice. Another breeding colony was established by crossbreeding male MFS (*Fbn1^C1041G/+^*) and female wildtype (*C57BL/6J*) mice to produce equal numbers of MFS and wildtype mice for the study. To determine and confirm mouse genotype, mouse tail samples were sent to Transnetyx (Transnetyx, Cordova, TN) for verification by PCR. Following genotyping, mice were divided into experimental groups (N=7-10) as follows: wildtype control (CTRL), MFS, *Cav1* KO, and MFS/*Cav1* KO. Mice (4 per cage) were housed under standard animal room conditions (a 12-hour light and dark cycle, 25°C), and fed on a standard Teklad Global 18% protein rodent diet (Envigo, USA). All animal care, breeding, and surgical procedures were done according to guidelines set by the National Institute of Health and the Midwestern University Institutional Animal Care and Use Committee [approved IACUC protocols MWU-2635 & MWU-2636].

### Sample Size Calculations

Sample size calculations were performed for different sets of experiments. For small chamber myography, the calculation was based on detecting a 20% decrease in aortic root diameter (lumen size) that corresponds to the threshold for surgical intervention in MFS aneurysm patients. The same parameters were used in our previous published reports using the same transgenic animal model of MFS. Based on our previous data, we performed a power analysis using G*Power 3.1, with an alpha level of 0.05 and a power of 0.90, resulting in a required sample size of minimum of 7-8 mice per group. For histological staining of elastin, we performed the calculation based on detecting a 20% decrease in elastin fragment counts within the aortic wall using our previously published data. The G*Power calculation, with an alpha level of 0.05 and a power of 0.90, resulted in a required sample size of minimum of 4 mice per group.

### Aortic Tissue Preparation

At 9 months of age, mice from all experimental groups were euthanized with 5% isoflurane and 2L/min of 100% oxygen inhalant followed by cervical dislocation. The thoracic cage was collected and placed in ice-cold aerated (95% O2-5% CO2) HEPES-PSS. Later, the whole length of thoracic aorta was dissected, cleaned of fat, and cut into 2mm segments (rings), with special care to protect the vessel’s endothelial layer. The ascending aorta (from root to arch) was also collected and fixed in 10% formalin buffer for histological processing and staining.

### Buffers and Reagents

Aortic segments were dissected, cleaned, and incubated in HEPES Physiological Salt Solution (HEPES-PSS) buffer (pH 7.43) containing 10 mM HEPES, 6 mM glucose, 1.8 mM CaCl_2_, 130 mM NaCl, 4 mM KCl, 4 mM NaHCO_3_ 1.2 mM MgSO_4_, 1.18 mM KH_2_PO_4_, and 0.03 mM EDTA, to simulate physiological conditions. The high KCl (K^+^) buffer (pH 7.43) contained 65mM NaCl, 80mM KCl, 1mM MgCl_2_, 10mM glucose, 5mM HEPES, 1.5mM CaCl_2_, and was used to assess smooth muscle viability and contractile ability. All pharmacological agonists and blockers including vasoconstricting agent phenylephrine (PE), vasodilatory agent acetylcholine (ACh), non-specific and reversible nitric oxide synthase (NOS) inhibitor L-N^G^-Nitro arginine methyl ester (L-NAME), and specific and potent inhibitor of inducible nitic oxide synthase (iNOS) 1400W dihydrochloride, were purchased from Sigma Millipore (Sigma Millipore, St. Louis, MO). All chemicals used in HEPES-PSS buffer were also purchased from Sigma Millipore.

### Transmission Electron Microscopy

We used transmission electron microscopy (TEM) to visualize caveolae structure in the mouse aortic wall. Aortic tissues were harvested from experimental mice, fixed in 2.5% glutaraldehyde in 0.1 M phosphate buffer (pH 7.4) for 24 hours at 4°C, and post-fixed in 1% osmium tetroxide. Tissues were then dehydrated through a graded ethanol series and embedded in epoxy resin. Ultrathin sections (70-90 nm) were cut using an ultramicrotome, collected on copper grids, and stained with uranyl acetate and lead citrate. TEM images were captured using a Philips CM 12 electron microscope operating at 80 kV.

### Measurements of aortic contraction and vasorelaxation

From each experimental mouse, four aortic segments (2mm in length) were mounted to a 4-unit chamber myograph for *ex vivo* study (DMT-USA Inc., Ann Arbor, MI). Force readings were generated and recorded via LabChart software (ADinstruments, Colorado Springs, CO). Segments were rested for 30min in continuously aerated 37°C HEPES-PSS (gassed continuously with 95% O_2_ and 5% CO_2_), and then stretched to their optimal tension of 6mN, as established by previous studies for C57BL/6 wild-type and MFS mice ^42^. Aortic segment viability was determined by challenging with 60mM KCl buffer twice, with 15 minutes of rest between each challenge. To assess smooth muscle contractility, segments were challenged with increasing doses (1 nM-50 µM) of phenylephrine (PE), the alpha-1-adrenergic vasoconstrictor agent. To assess endothelial function, pre-constricted segments were subjected to increasing doses (50 pM- 1 µM) of acetylcholine (ACh). Aortic relaxation data are expressed as a percentage of the pre-constricted aortic diameters in each experiment. To evaluate the effects of NO, segments were treated with L-arginine methyl ester hydrochloride (L-NAME), a non-selective and general nitric oxide synthase inhibitor. To evaluate the effects of NO originated from inducible NOS (iNOS) activation, segments were treated with 1400W, an irreversible and specific iNOS inhibitor.

### Measurements of aortic diameters and rupture points

To assess the internal lumen diameter of mouse aorta, 2-mm aortic sections from the region adjacent to the aneurysm area were harvested and anchored onto the myograph chambers between two pins as described previously ^42^. In the myograph chamber, the distance between the two pins (on which the aortic ring is anchored) at the initial point, where the generated force is first recorded (when the aortic wall initially touches the pin), is an indicator of the aortic luminal diameter in the resting condition. In another set of experiments, we tested the wall strength of the aortic segments by gradually increasing the distance between the two myograph pins (200 µm diameter) in 100 µm intervals, which correlates with an increase in the estimated length of vascular smooth muscle within the vessel wall. The stretching protocol was repeated until the aortic segment could not maintain its tone, and the force (stress point) at which the anchored aortic segment ruptured was recorded as the rupture point, as previously described ^42^.

### Aortic wall elastin staining

The aortic arch of each mouse was collected, carefully cleaned, and fixed in 10% buffered formalin for 24 hours and then transferred into 70% ethanol to be used for histological studies. Samples were sent to HistoWiz (Brooklyn, NY, USA) for processing, embedding, sectioning, and Verhoeff-van Gieson staining. To ensure consistent imaging, all stained slides were imaged, using an Olympus light microscope and Zeiss AXIO digital camera, by an independent investigator blinded to genotype. Verhoeff-van Gieson staining was used to analyze medial elastin structure in serial cross sections (10µm) of the aortic arch after rehydration of tissue sections. Elastin fragmentation within the aortic wall sections was examined using ImageJ (Media Cybernetics, Bethesda, MA) object tracing tools by tracing elastin fibers measured in pixels and converted to micrometers. Processing, staining, and analysis was blinded to prevent bias.

### Statistical Analysis

Small vessel chamber myography experiments and data analysis were performed by a single investigator blinded to animal genotypes and treatment. Histological staining and images were processed by HistoWiz (Brooklyn, NY, USA) blinded to genotype and treatment. Image and data analysis were also performed by an independent lab member blinded to animal genotypes and treatment. Statistical analysis and construction of all concentration-response curves and graphs were performed using GraphPad Prism 10.2.3 Software. A two-way analysis of variance (ANOVA) test was used to compare three or more groups, followed by a *post-hoc Tukey* test for multiple statistical comparisons. Data are presented as Mean ± SE, where a *P* value less than 0.05 (*P* < 0.05) is considered as significant.

## RESULTS

### Genetic Cav1 deletion disrupts the formation of caveolae invagination in the aortic wall

In addition to confirming the deletion of the *Cav1* gene in mice using PCR, we verified that the ablation of *Cav1* in both control and MFS mice had the anticipated effects on the structure of caveolae in the aortic wall. Through observational and qualitative assessments via electron microscopy (EM) images, we found that the deletion of *Cav1* disrupted the formation of caveolae invaginations on the cell membrane (**Fig. 1**). In CTRL mice aorta, abundant and well-formed caveolae invaginations are clearly visible (white arrows), characterized by flask-shaped membrane invaginations approximately 50-100 nm in diameter (**Fig. 1A, white arrows**). Aortic sections isolated from MFS mice display a reduction in the number of open invaginations (caveolae) on the membrane of endothelial cells (**Fig. 1B**). Likewise, aortic rings from *Cav1* knockout mice (CTRL/*Cav1KO* & MFS/*Cav1KO*) exhibit a reduction in the number of visible caveolae invaginations on the membrane surface (**Fig. 1C & 1D**). The plasma membrane appeared mostly smooth and devoid of the characteristic invaginated structures seen in the CTRL mice aortic sections. This disruption suggests that Cav1 protein plays a critical role in the biogenesis and maintenance of caveolae on cell membranes, and the absence of Cav1 likely impairs the assembly of all associated proteins necessary for caveolae invagination on the membrane surface.

**Figure 1.**
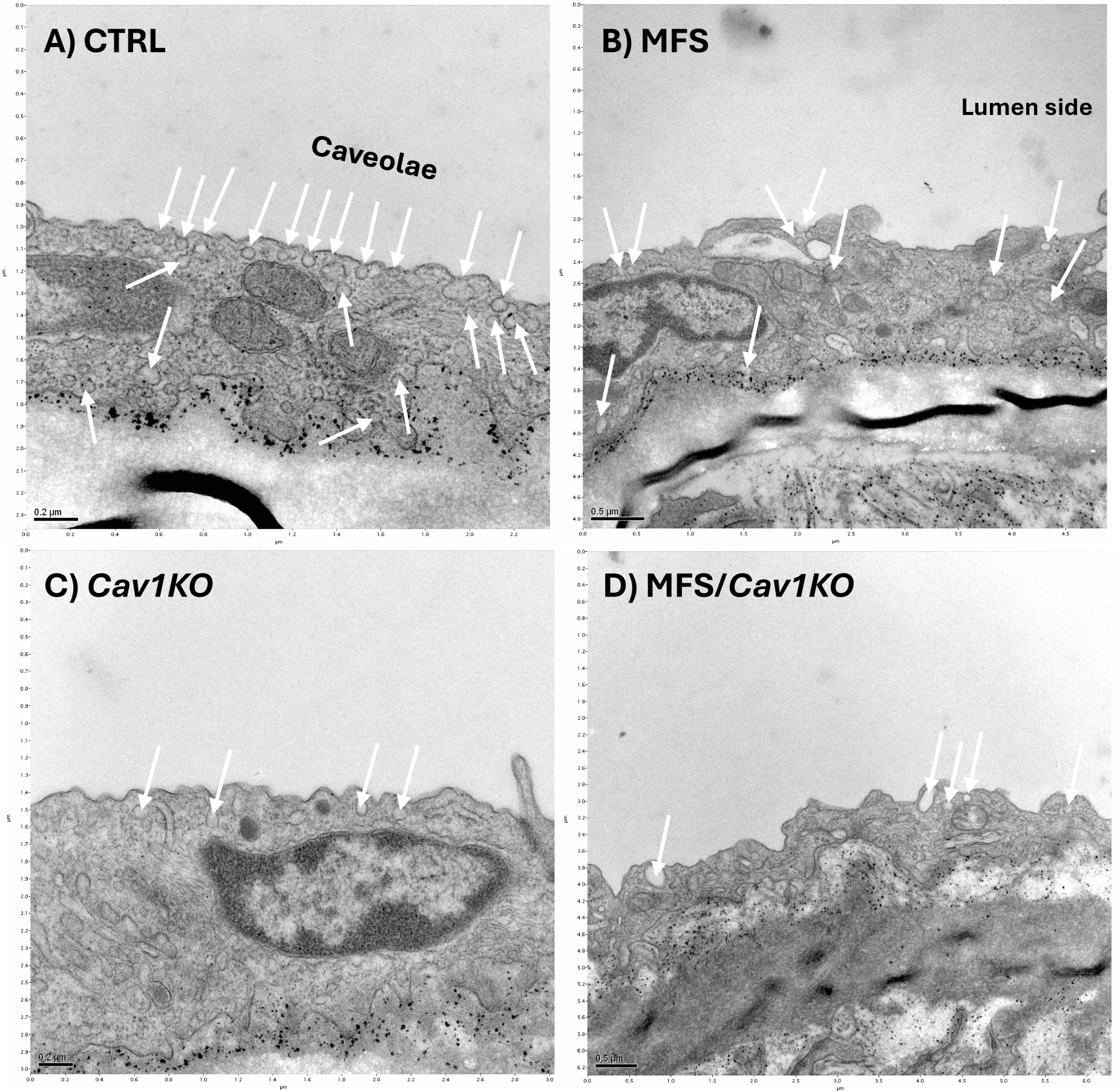
Genetic *Cav1* deletion disrupts the formation of caveolae invagination in the aortic wall. Representative transition electron microscopy images of 9-month-old aortic sections in CTRL **(A)**, MFS **(B)**, *Cav1KO* **(*C*)**, and MFS/*Cav1KO* **(D)** mice visualize the structure of caveolae invaginations in the aortic wall and shows that deletion of *Cav1* gene impacts the structural integrity and membrane distribution of caveolae.

### Deletion of Cav1 gene doesn’t affect the progression of aortic root growth in MFS mice

To determine the impact of *Cav1* deletion on aortic root growth and aneurysm progression in male and female MFS mice, we measured the internal diameters of aortic rings isolated from mice. As expected, aortic root diameters are significantly increased in both male and female MFS mice as compared to healthy CTRL with no sex differences observed (**Fig. 2A**). Interestingly, *Cav1* deletion had no effects on aortic root growth and aneurysm formation in either male or female MFS/*Cav1KO* groups (**Fig. 2B**).

**Figure 2.**
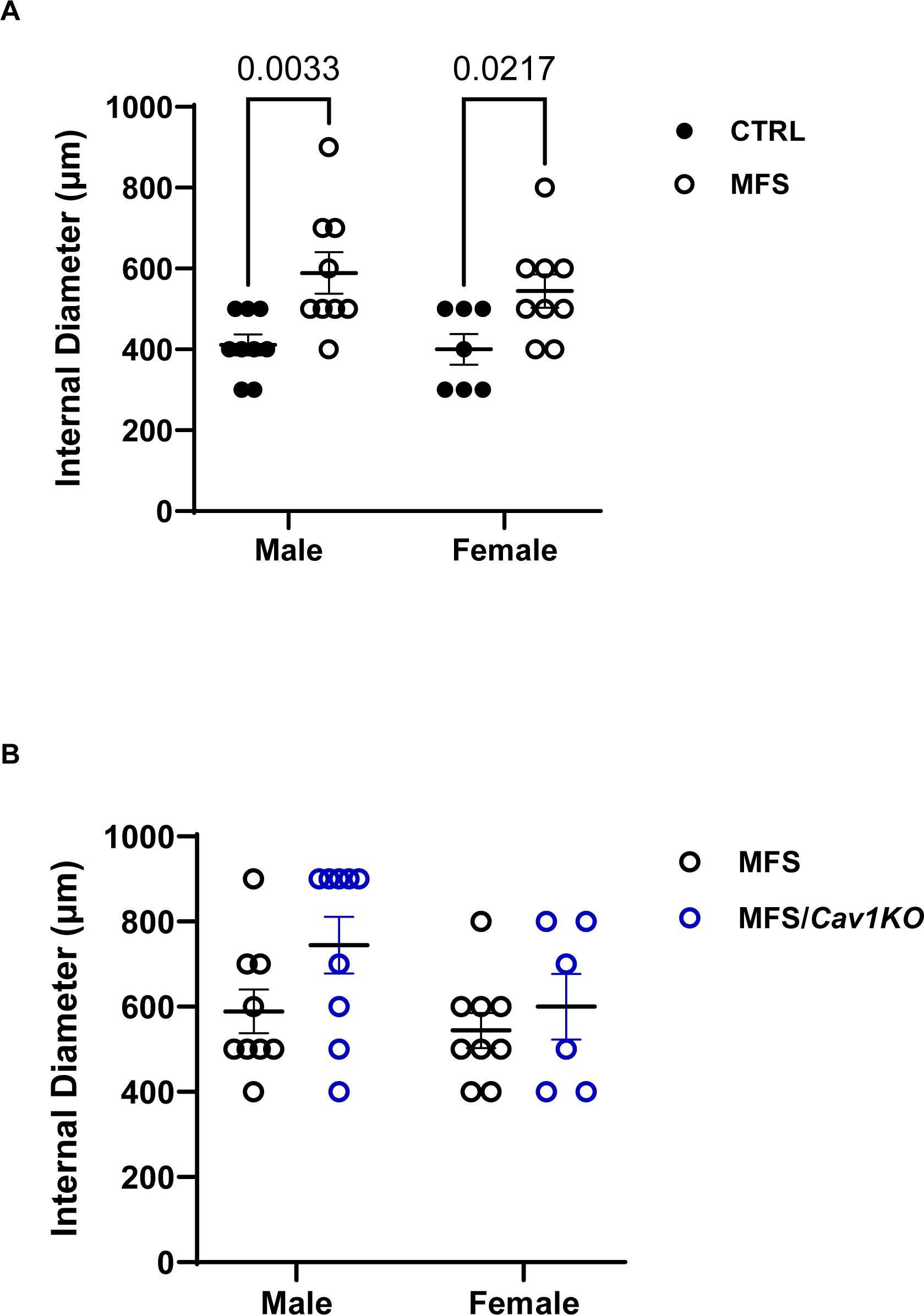
Deletion of *Cav1* gene doesn’t affect the progression of aortic root growth in MFS mice. Presented bargraphs showcase measurements of aortic root diameter using the myograph chambers. **A)** Aortic root diameters are increased in both male and female MFS mice compared to age- and sex-matched healthy CTRL mice. **B)** *Cav1* gene deletion has no impact on artic root growth in both male and female MFS mice aorta. (Means ± SE, N = 6-9 mic/group, Two Way ANOVA followed by Tukey’s pairwise comparison, *P* ≤ 0.05).

### Deletion of Cav1 increases endothelium-dependent vasorelaxation in MFS mice

Given the well-documented inhibitory effect of Cav1 protein on endothelial NO production, we assessed the impact of *Cav1* deletion on endothelium-dependent vasorelaxation in male and female CTRL and MFS mice. In myograph chambers, aortic segments were pre-constricted with the sub-maximum concentration of PE (10 µM), followed by the application of the vasorelaxant agent, acetylcholine (Ach), in a dose-dependent manner (**Supplementary Fig. S1**). We observe a significant decrease in maximum Ach-induced aortic relaxation in male MFS mice at 9 months of age (**Fig. 3A**), with no difference detected between age-matched female CTRL and MFS mice (**Fig. 3A**). It is noteworthy that aortic relaxation is markedly higher in female MFS mice compared to age-matched male MFS mice (**Fig. 3A**). Deletion of Cav1 resulted in a significant increase in Ach-induced relaxation in both male and female MFS/*Cav1KO* mice compared to age and sex-matched MFS groups, indicating that *Cav1* deletion leads to an increase in NO production in MFS mice aortic rings (**Fig. 3B**).

**Figure 3.**
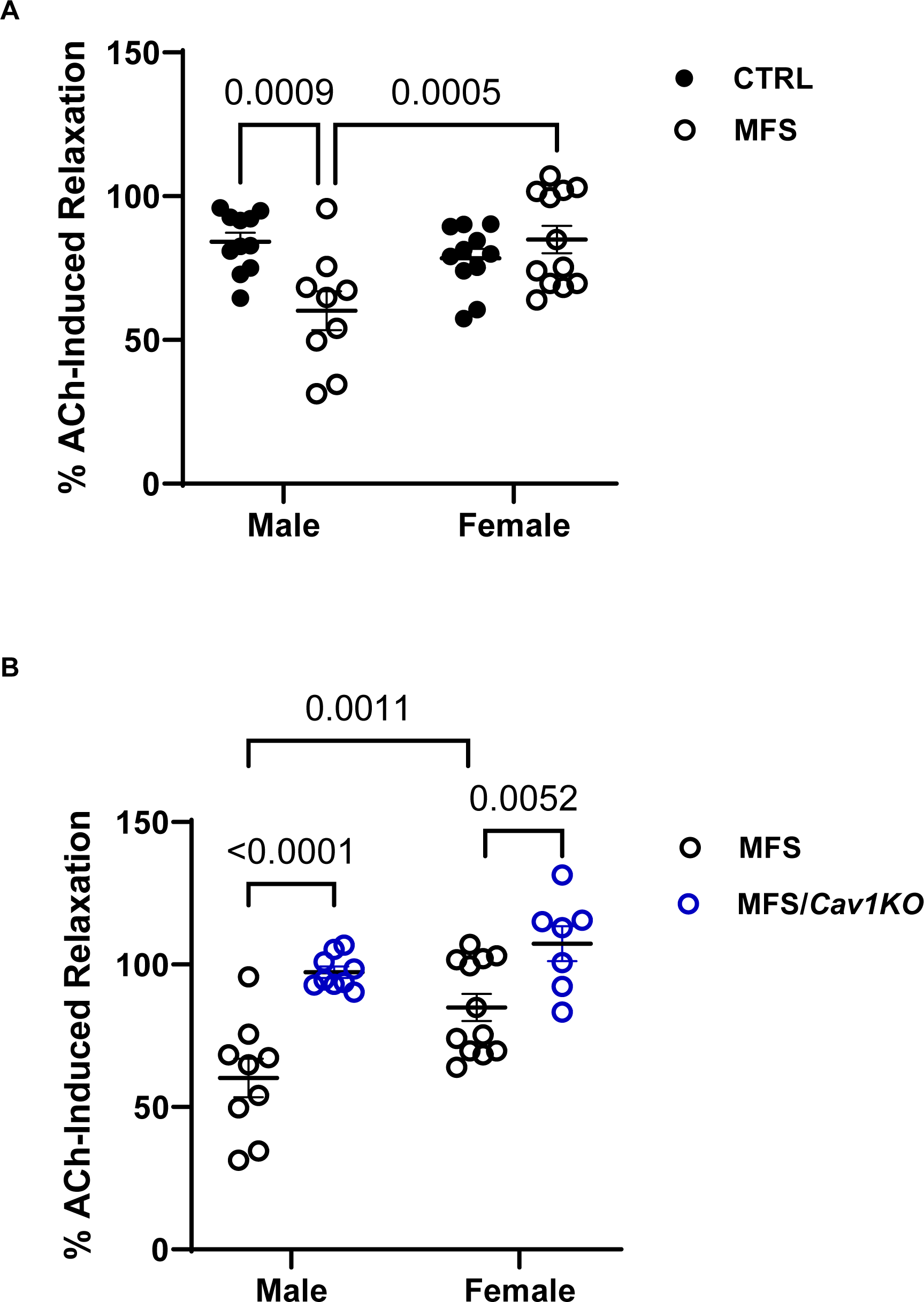
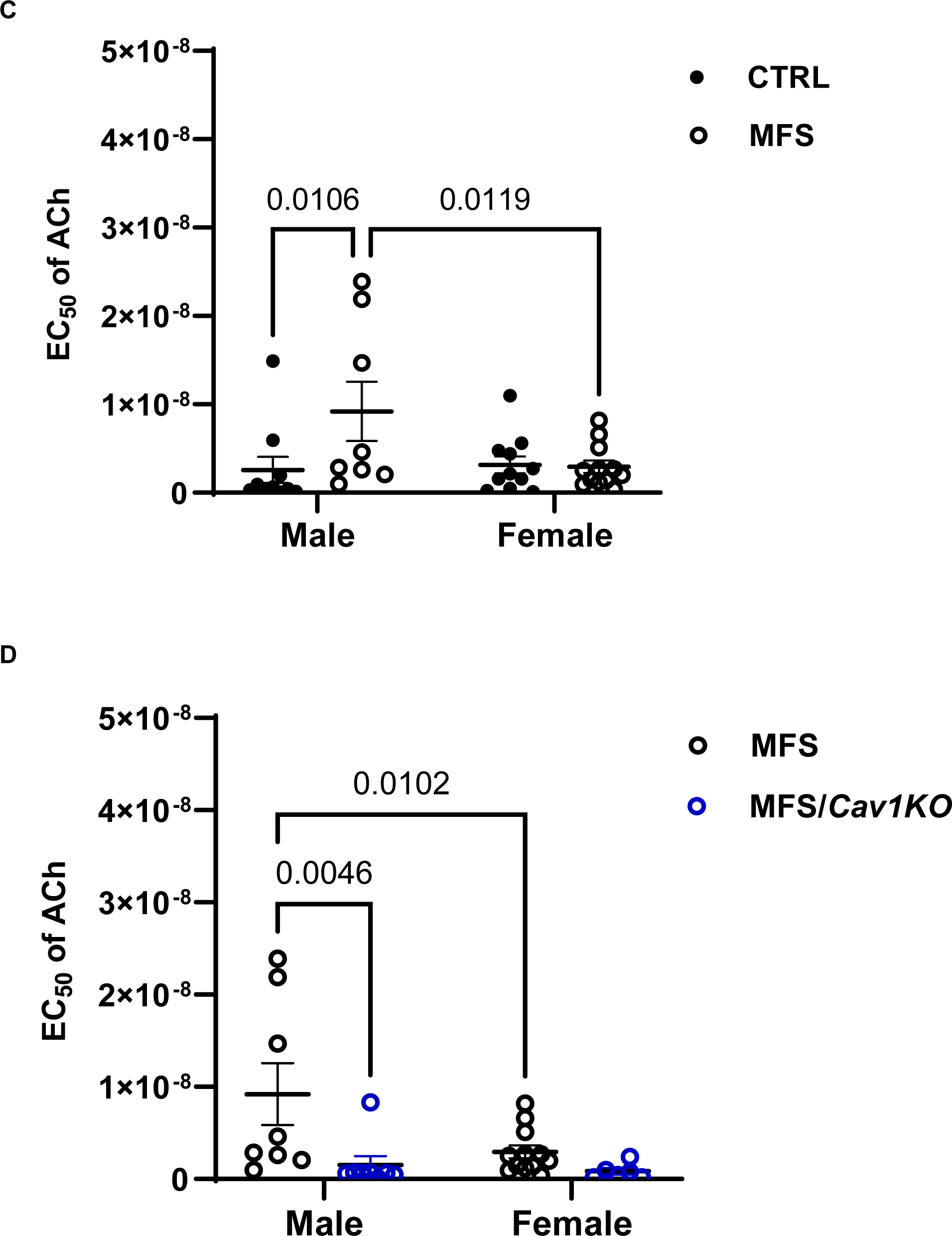
Deletion of *Cav1* increases endothelium-dependent vasorelaxation in MFS mice. Bargraphs present acetylcholine (Ach)-induced aortic relaxation of aortic rings isolated from different experimental groups using isometric small chamber myography. **A)** Aortic relaxation in response to Ach is significantly decreased in aortic rings isolated from male and female MFS mice compared to age- and sex-matched health CTRL aorta. **B)** Cav1 gene deletion leads to an increase in Ach-induced aortic vasorelaxation in male and female MFS/*Cav1KO* mice compared to age-and sex-matched MFS mice, indicating significant increases in aortic wall NO production in the absence of Cav1 protein. **C)** EC_50_ values for Ach is markedly increased in male MFS mice as compared to sex- and age-matched CTRL, but no difference is observed in age-matched female CTRL and MFS aorta. When we compare age-matched male and female CTRL mice, the EC_50_ values are lower in female aorta. **D)** Deletion of *Cav1* results in a significant drop in Ach EC_50_ values in age-matched male MFS/*Cav1KO* aorta as compared to MFS mice, indicating an increased sensitivity of aortic endothelial layer to Ach treatment in male mic aortic rings. However, in female mice, *Cav1* deletion doesn’t impact aortic endothelial cell sensitivity to Ach. (Means ± SE, N = 9-12 mic/group, Two Way ANOVA followed by Tukey’s pairwise comparison, P ≤ 0.05).

To investigate whether the observed increases in Ach-induced vasorelaxation in male and female MFS/*Cav1KO* mice aorta are due to changes in aortic endothelial cells sensitivity and response to NO, we calculated the EC_50_ values for Ach from the dose-response curves (**Supplementary Fig. S1**). Our data shows that the EC_50_ values for male MFS aorta is higher compared to male CTRL mice, with no difference observed between female CTRL and MFS aortic rings (**Fig. 3C**). We also observe a significant decrease in Ach EC_50_ values in female MFS aortic rings compared to age-matched male MFS aorta, indicating that aortic rings isolated from female MFS mice, are more sensitive to Ach (**Fig. 3C**). The deletion of *Cav1* causes further drops in EC_50_ values in male MFS/*Cav1KO* mice aorta, with no effects observed in female MFS/*Cav1KO* mice (**Fig. 3D**). These observations suggest that *Cav1* deletion-mediated increase in aortic relaxation in male MFS aorta could be partially attributed to enhanced sensitivity of aortic endothelial cells to the vasorelaxant agent acetylcholine. Interestingly, in healthy CTRL mice, while *Cav1* deletion causes an increase in aortic relaxation in female CTRL/*Cav1KO* mice, such effects were not detected in males CTRL/*Cav1KO* subjects, suggesting a sex-dependent effect in CTRL aorta (**Supplementary Fig. S2**).

### Cav1 gene deletion reduces phenylephrine-induced aortic contraction in CTRL & MFS mice

We evaluated the impact of *Cav1* deletion on smooth muscle membrane polarization and force generation, mediated by calcium influx through voltage-gated calcium channels (VGCC)) in a myograph chamber by subjecting isolated aortic rings from 9-month-old male and female CTRL, MFS, and MFS/*Cav1*KO mice to 70mM KCl buffer solution. In male mice, aortic contraction in response to KCl was not significantly different between CTRL and MFS groups (**Fig. 4A**). On the other hand, in female MFS mice, aortic contraction in response to KCl was significantly lower compared to CTRL groups (**Fig. 4A**). Deletion of the *Cav1* gene significantly reduced KCl-induced aortic contraction in both male and female MFS/*Cav1*KO groups, indicating Cav1’s crucial role in regulating the normal function of VGCC in aortic rings independent of sex (**Fig. 4B**).

**Figure 4.**
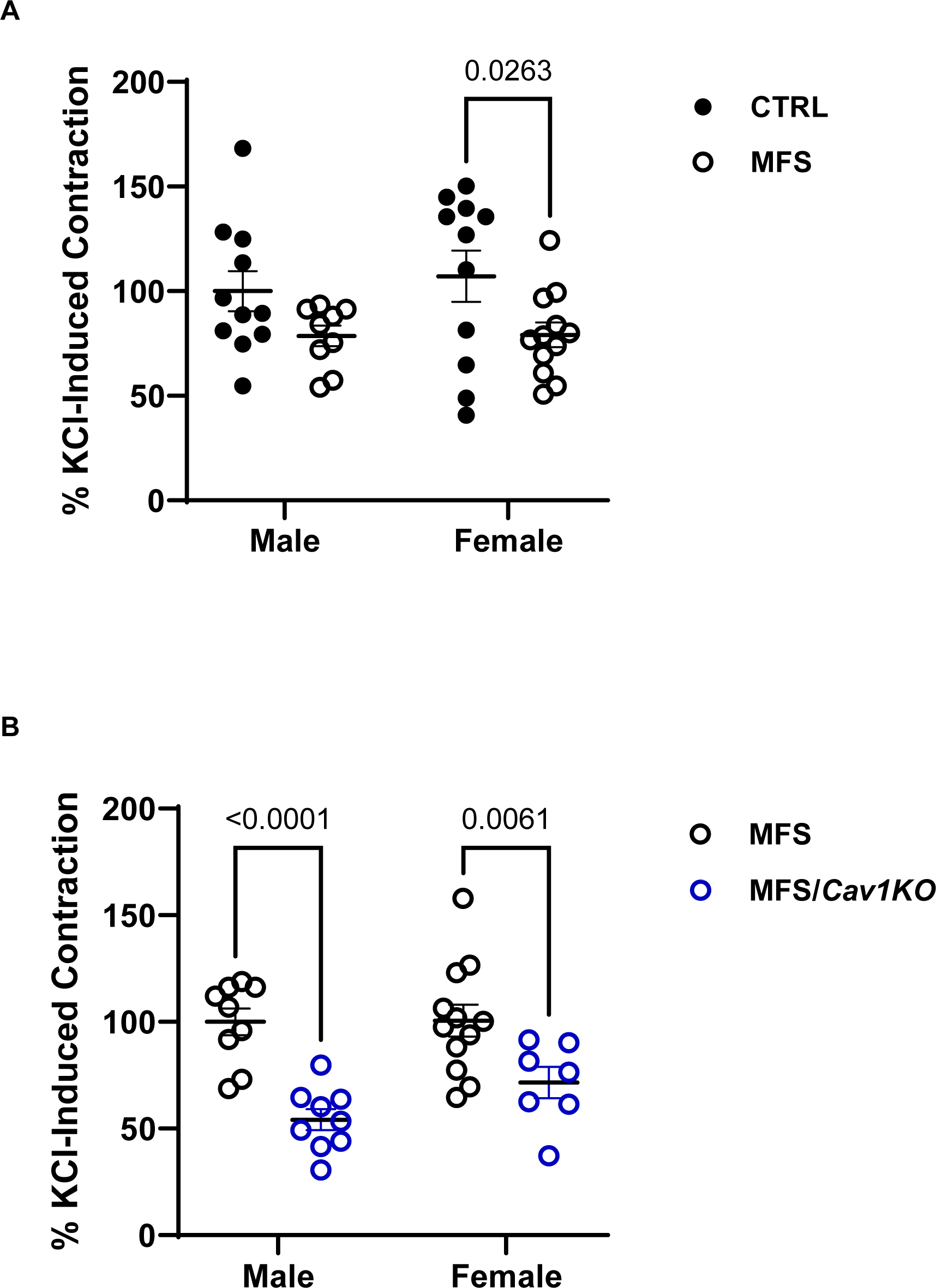
*Cav1* gene deletion reduces KCl-induced aortic contraction in MFS mice. Bargraphs present maximum force generation in isolated aortic rings in response to KCl (60mM) in the myograph chambers. **A)** KCl-induced aortic contraction in reduced in female MFS mice compared to age- and sex-matched controls. No genotype associated differences are observed in age-matched male CTRL and MFS. **B)** *Cav1* deletion reduces KCl-induced smooth muscle contraction in male and female MFS/*Cav1KO* mice compared to age- and sex-matched MFS aorta. (Means ± SE, N = 9-12 mic/group, Two Way ANOVA followed by Tukey’s pairwise comparison, P ≤ 0.05).

To further investigate the impact of *Cav1* deletion on receptor-mediated smooth muscle contraction, we subjected aortic rings isolated from all experimental groups to increasing doses (10 nM-50 µM) of the vasoconstricting agonist phenylephrine (PE) in myograph chambers (**Supplementary Fig. S3**). Our findings show that at 9 months of age, the maximum PE-induced contractile forces (E_max_) are significantly reduced in both male and female MFS mice compared to age and sex-matched CTRL subjects (**Fig. 5A**), with female MFS aorta showing lower levels of contraction compared to male MFS aorta (**Fig. 5B**). Ablation of the *Cav1* gene causes further drops in PE-induced contraction in male, but not in female MFS/*Cav1KO* aorta (**Fig. 5B**), indicating a sex-dependent effect.

**Figure 5.**
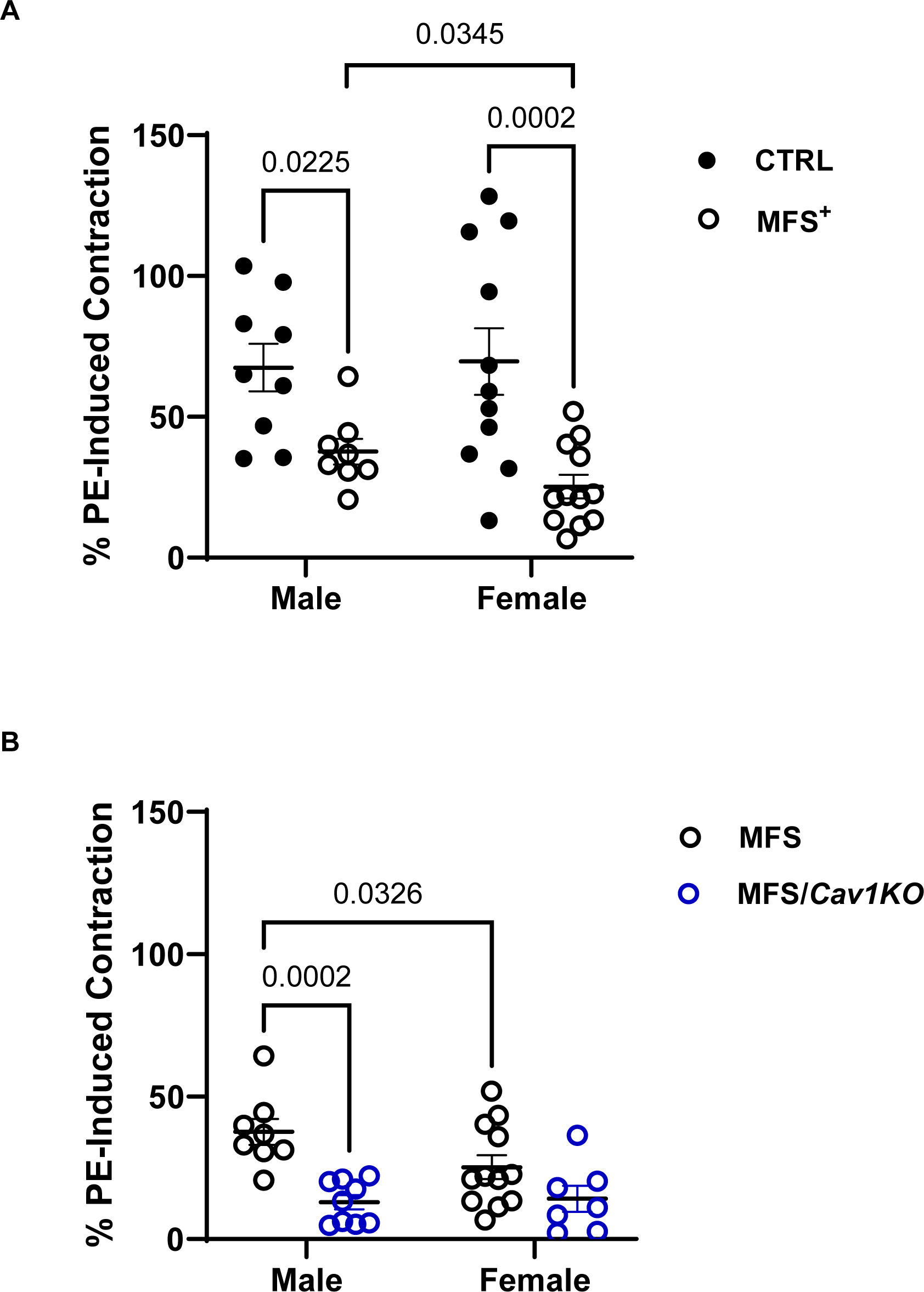
*Cav1* gene deletion decreases phenylephrine-induced aortic contraction in CTRL & MFS mice. Bargraphs present mice aortic rings contractions in response to sub-max concentration (1 µM) of phenylephrine (PE) in the myograph chambers. **A)** In both male and female MFS mice, aortic rings contractions in response to PE is significantly lower compared to healthy CTRL aorta. When we compare age-matched male and female MFS aorta, we observed a significant reduction in female MFS aortic contraction as compared to male MFS mice. **B)** *Cav1* deletion reduces PE- induced aortic contraction in male MFS mice, but not in females, indicating a sex-specific effect on aortic wall contraction. However, it’s important to notice that *Cav1* deletion reduces PE-induced contraction in male MFS aorta to levels that are not different from the values observed in female MFS/*Cav1KO* mice. (Means ± SE, N = 7-10 mic/group, Two Way ANOVA followed by Tukey’s pairwise comparison, P ≤ 0.05).

### Cav1 gene deletion reduces mouse aortic contraction by increasing NO production

To investigate whether the *Cav1KO*-mediated decrease in aortic contraction is facilitated through an increase in the NO production, aortic segments were pre-incubated with a non-selective nitric oxide synthase (NOS) inhibitor, L-NAME (200 µM) in the myograph chambers for 30 minutes before the application of PE. Pre-treatment of aortic rings with L-NAME causes significant increases in PE-induced vasocontraction peaks in both male (**Fig. 6A**) and female (**Fig. 6B**) CTRL and MFS mice, due to an L-NAME-mediated decrease in the endogenous NO production within the aortic walls, with no differences observed between sexes (data not shown). Furthermore, in both male (**Fig. 6C**) and female (**Fig. 6D**) MFS/*Cav1KO* mice, pre-treatment of aortic rings with L-NAME induces further increases in PE-induced contraction forces, indicating that *Cav1* deletion causes an even more significant increases in endogenous NO production in MFS mouse aortic rings, further highlighting the notion that Cav1 regulates MFS mouse aortic contraction through NO-dependent mechanisms.

**Figure 6.**
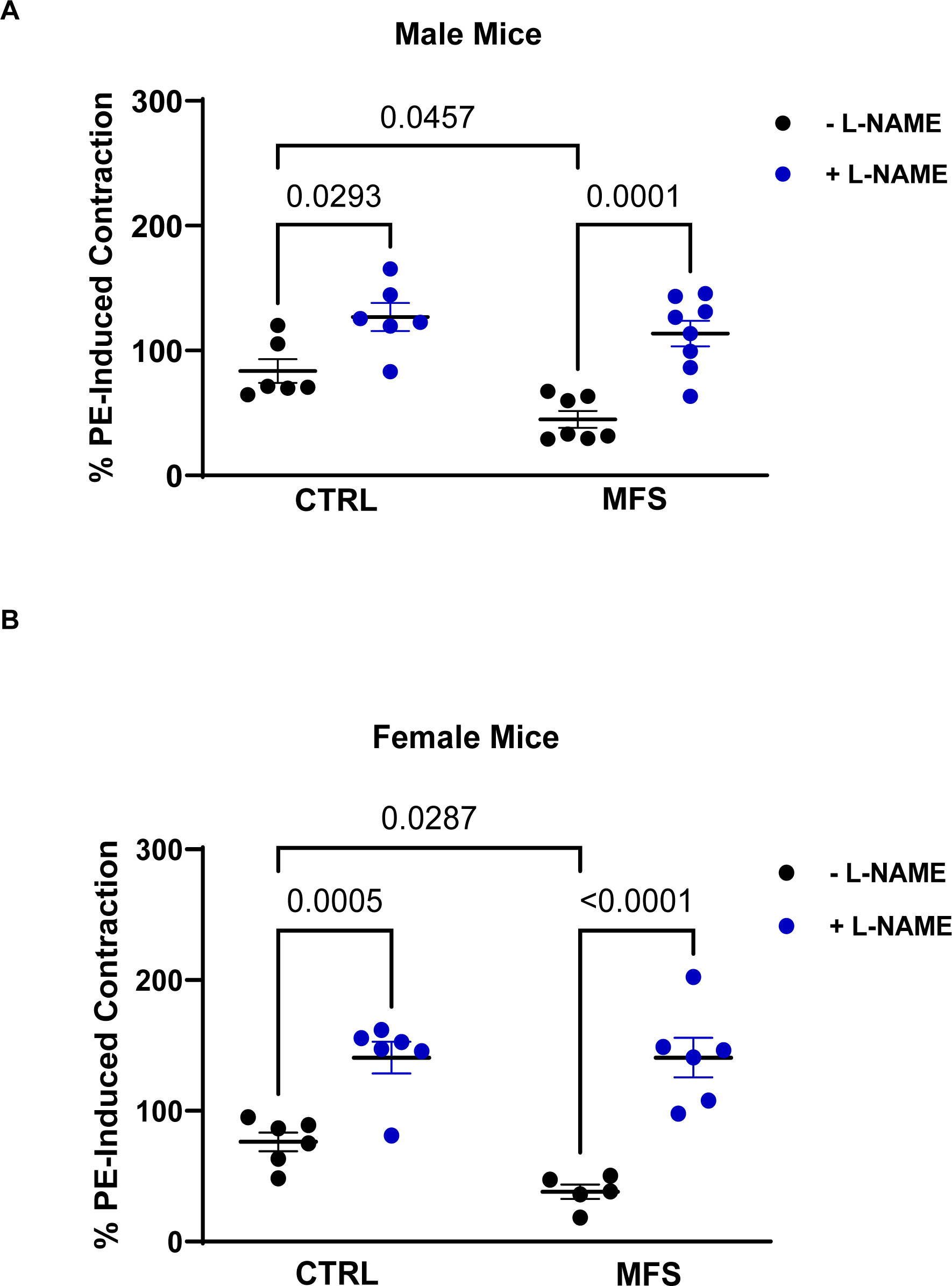
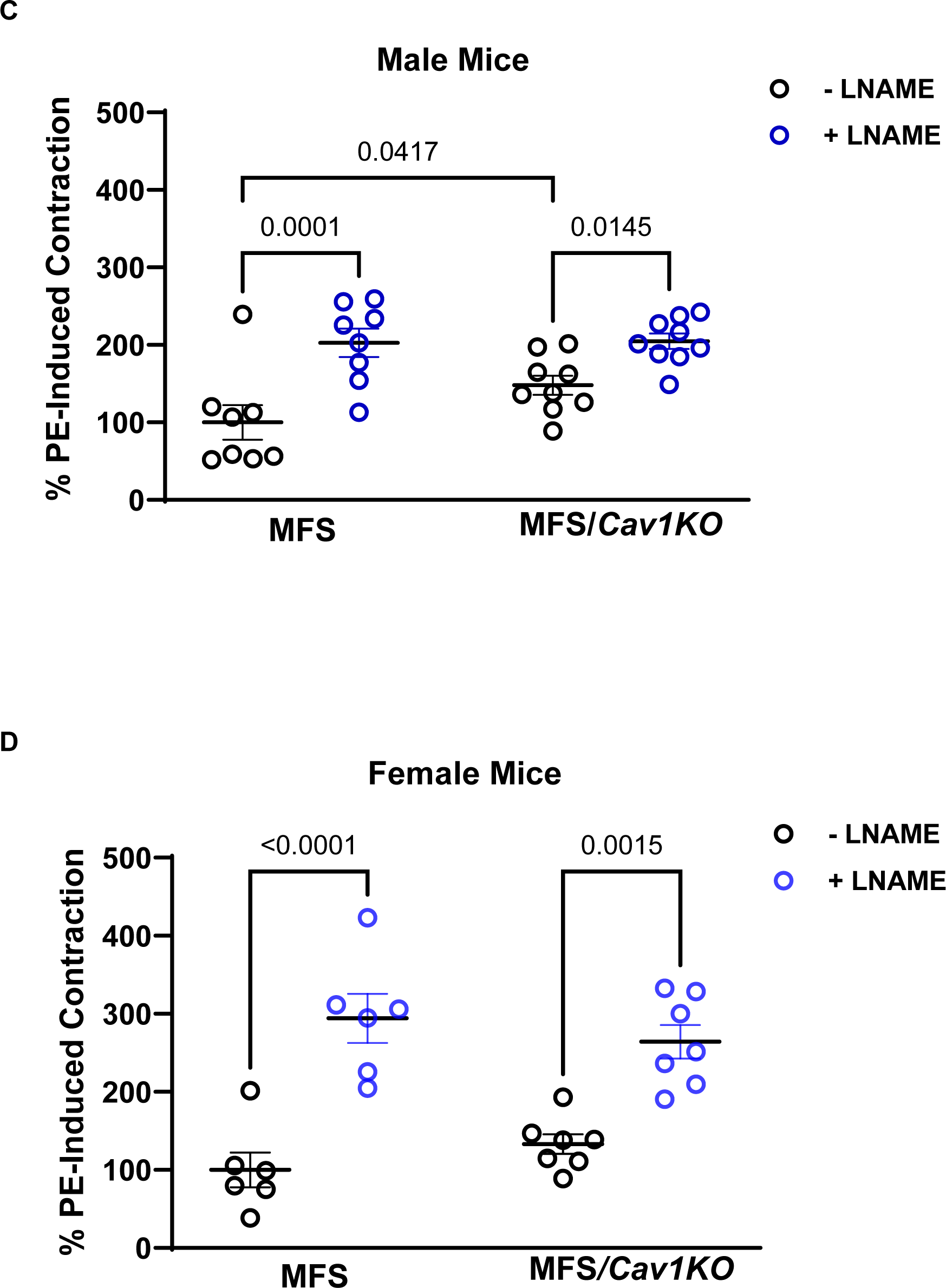
*Cav1* gene deletion decreases mouse aortic contraction by increasing NO production. Bargraphs present aortic contraction in response to sub-maximum concentration (1 µM) of phenylephrine (PE) in CTRL, MFS, and MFS/*Cav1KO* mice in the absence or presence of L-NAME (200 µM), which is a general inhibitor of NO synthesis. **A)** Pre-treatment of aortic rings with L-NAME increases aortic root contraction in age-matched male CTRL and MFS aorta. **B)** Pre-treatment of aortic rings with L-NAME also increases aortic root contraction in age-matched female CTRL and MFS aorta. **C)** Similar to male MFS mice, in male MFS/Cav1KO mice aorta, pre-treatment of aortic rings with L-NAME significantly increases aortic wall contraction in response to PE. **D)** Similar to the phenomenon observed in female MFS mice aorta, L-NAME increases PE-induced aortic contraction in female MFS/*Cav1KO* mice. (Means ± SE, N = 6-9 mic/group, Two Way ANOVA followed by Tukey’s pairwise comparison, P ≤ 0.05).

### Cav1 deletion-mediated increase in aortic NO production is not through iNOS

The production of NO in the vascular system is controlled by three different isoforms of NOS including endothelial nitric oxide synthase (eNOS), neuronal NOS (nNOS), and inducible NOS (iNOS). Among these three isoforms, iNOS is more involved in inflammatory responses.

Previous studies in aortic tissue from the MFS mouse model and the plasma isolated from MFS aneurysm patients have shown an increase in iNOS expression ^43–46^. These reports suggest the possibility that *Cav1* gene deletion may increase the expression and activity of iNOS as a potential source for increased endogenous NO in MFS/*Cav1*KO mice. To narrow down the source for increased NO production, we utilized a potent and specific inhibitor of iNOS, N-3- Aminomethyl benzyl acetamidine (1400W) in the myograph chamber. In our experiments, aortic rings were pre-incubated with 1400W (1 µM) for 30 minutes prior to PE application. Pre-treatment of aortic rings with the iNOS inhibitor did not impact the *Cav1*KO-mediated decrease in aortic contraction in either male or female MFS/*Cav1*KO aortic rings (**Fig. 7**). This important observation implies that iNOS expression is not affected by the *Cav1* gene deletion in MFS mice. Consequently, this raises the possibility that eNOS and/or nNOS are affected by *Cav1* gene deletion. Interestingly, pre-treatment of MFS mouse aortic rings with 1400W increases PE-induced contraction, which is in line with previous reports of an increase in iNOS activity in MFS aortic tissue ^43–46^.

**Figure 7.**
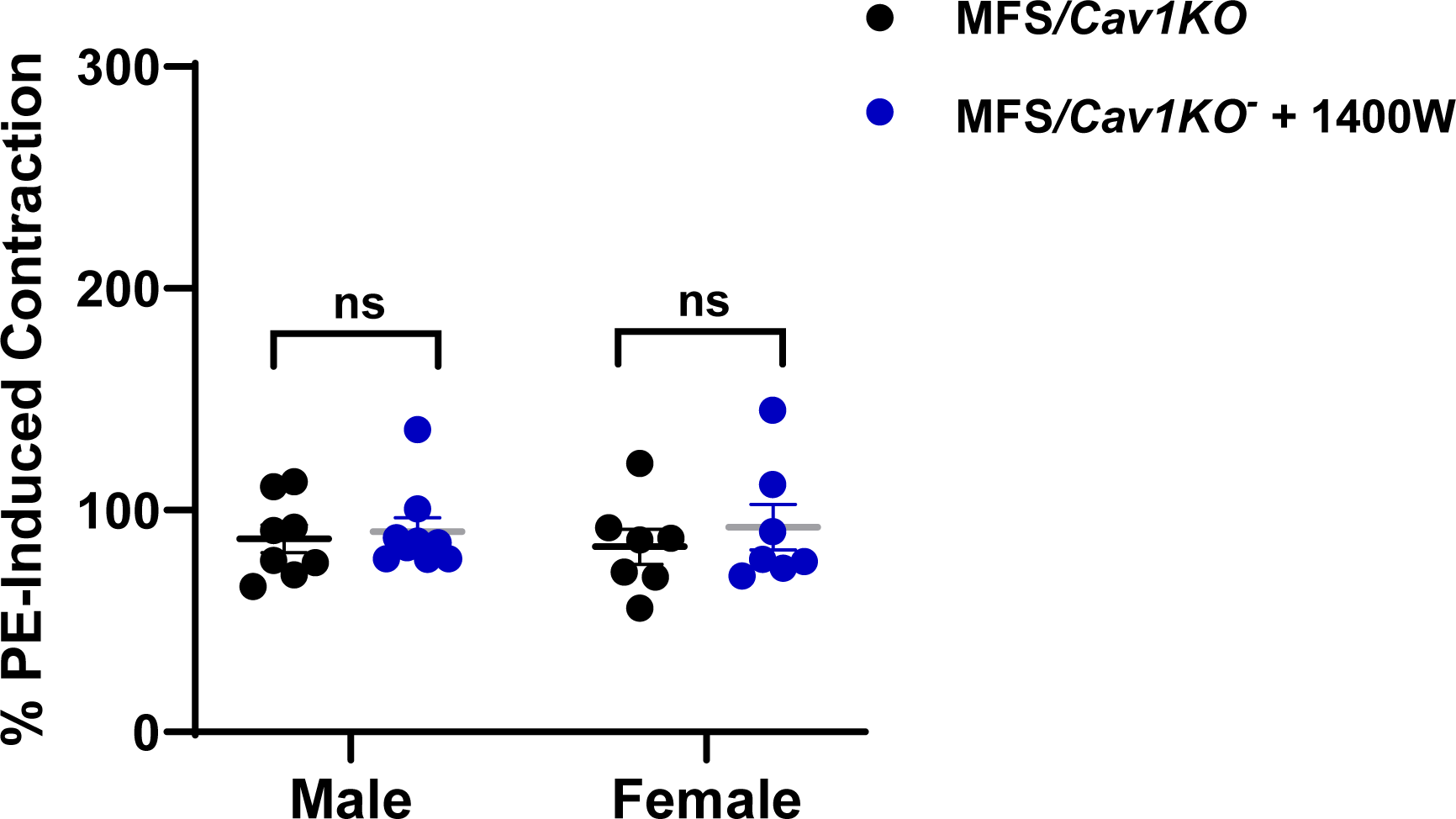
*Cav1* deletion-mediated increase in aortic NO production is not through iNOS. Presented bargraphs showcase the impact of 1400W, a potent inhibitor of inducible NOS (iNOS) on PE-induced contraction in aortic rings isolated from male and female MFS/*Cav1KO* mice. Data clearly shows that pre-treatment of aortic rings with 1400W (1 µM) has no effects on PE- induced aortic contraction in age-matched male and female mice, indicating that the excessive increase in aortic endogenous NO production in the absence of Cav1 is not mediated by iNOS activation (Means ± SE, N = 6-8 mic/group, Two Way ANOVA followed by Tukey’s pairwise comparison, P ≤ 0.05).

### Aortic wall strength and elastin structure are improved in male MFS/Cav1KO mice

It is well established that MFS aneurysm is associated with aortic wall weakening and loss of elasticity. To determine the effects of the *Cav1* gene deletion on aortic wall strength, the aortic segment rupture points were assessed at 9 months of age in all experimental groups using small vessel chamber isometric myography. The rupture point of aortic segments represents the maximum force generated by each segment at the point of maximum stretch, just before the aortic wall ruptures. Based on our findings, both male and female MFS mice demonstrated decreased aortic wall strength (reduced force generation at the rupture point) compared to age- and sex-matched CTRL mice (**Fig. 8A**). When we compare CTRL and CTRL/*Cav1KO* groups, only male, but not female CTRL/*Cav1KO* mice show an improvement in maximum wall strength (**Fig. 8B**). Similarly, when MFS aortic rings are compared with MFS/*Cav1KO* aorta, only male MFS/*Cav1KO* show improvements in aortic wall maximum strength (**Fig. 8C**). These findings suggest that the effects of *Cav1* deletion on aortic wall strength and structural integrity in both CTRL and MFS mice are sex-dependent.

**Figure 8.**
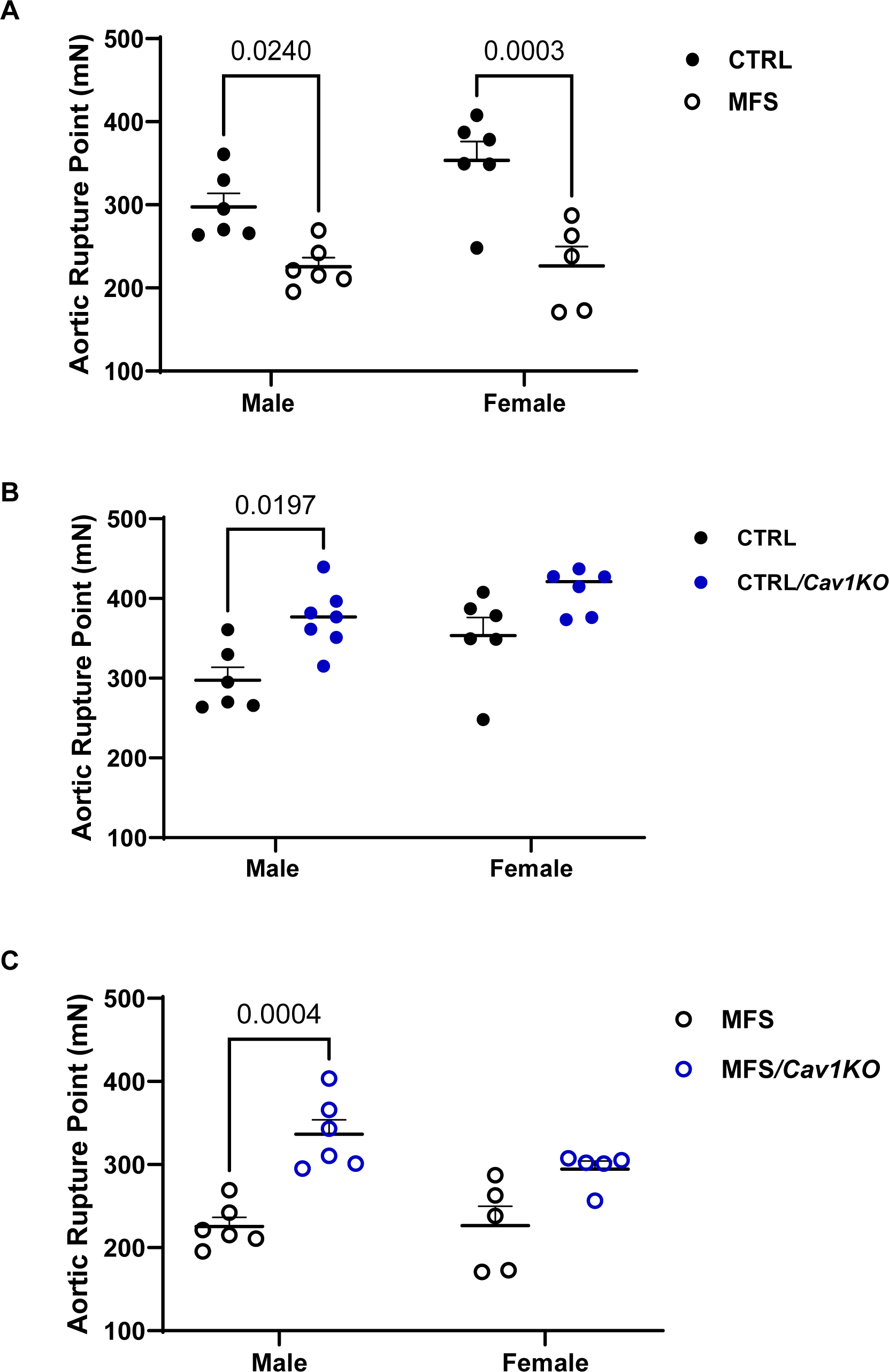
Aortic wall strength is markedly improved in male MFS/*Cav1KO* mice. Scatter plot graphs show the force generated (mN) at the rupture point. The rupture point of these aortic segments represents the maximum force generated by each segment at the point of maximum stretch, just before the aortic wall ruptures. **A)** As expected, aortic wall rupture point is significantly reduced in both male and female MFS groups as compared to age- and sex-matched healthy CTRL mice. **B)** Deletion of *Cav1* gene increases the aortic rings rupture points in male CTRL mice, with not effects observed in female CTRL subjects. **C)** Similarly, *Cav1* deletion further increases aortic wall strength (rupture point) only in male MFS mice, indicating a sex-dependent effect. (Means ± SE, N = 4 mic/group, Two Way ANOVA followed by Tukey’s pairwise comparison, P ≤ 0.05).

In MFS, aortic root aneurysm is associated with aortic wall medial elastin fragmentation. Since the *Cav1* deletion only improved male MFS mice aortic wall strength, we did a follow up study to determine the effects of the *Cav1* deletion on aortic wall elastin structural integrity in male MFS mice using van Gieson staining, where elastin fibers are stained in dark purple in the medial layer of the aortic wall (**Fig. 9A**). As previously demonstrated, male MFS mice show increased elastin fragmentation (**Fig. 9B**) and decreased elastin fiber length (**Fig. 9C**) in the aortic wall compared to male CTRL mice. In addition, *Cav1* deletion significantly improves aortic elastin structure in male MFS/*Cav1KO* mice, as evidenced by decreased elastin fragments count (**Fig. 9B**) and increased elastin fibers length (**Fig. 9C**) in the aortic wall. Collectively, these findings indicate improved aortic wall elasticity in male MFS/*Cav1KO* mice.

**Figure 9.**
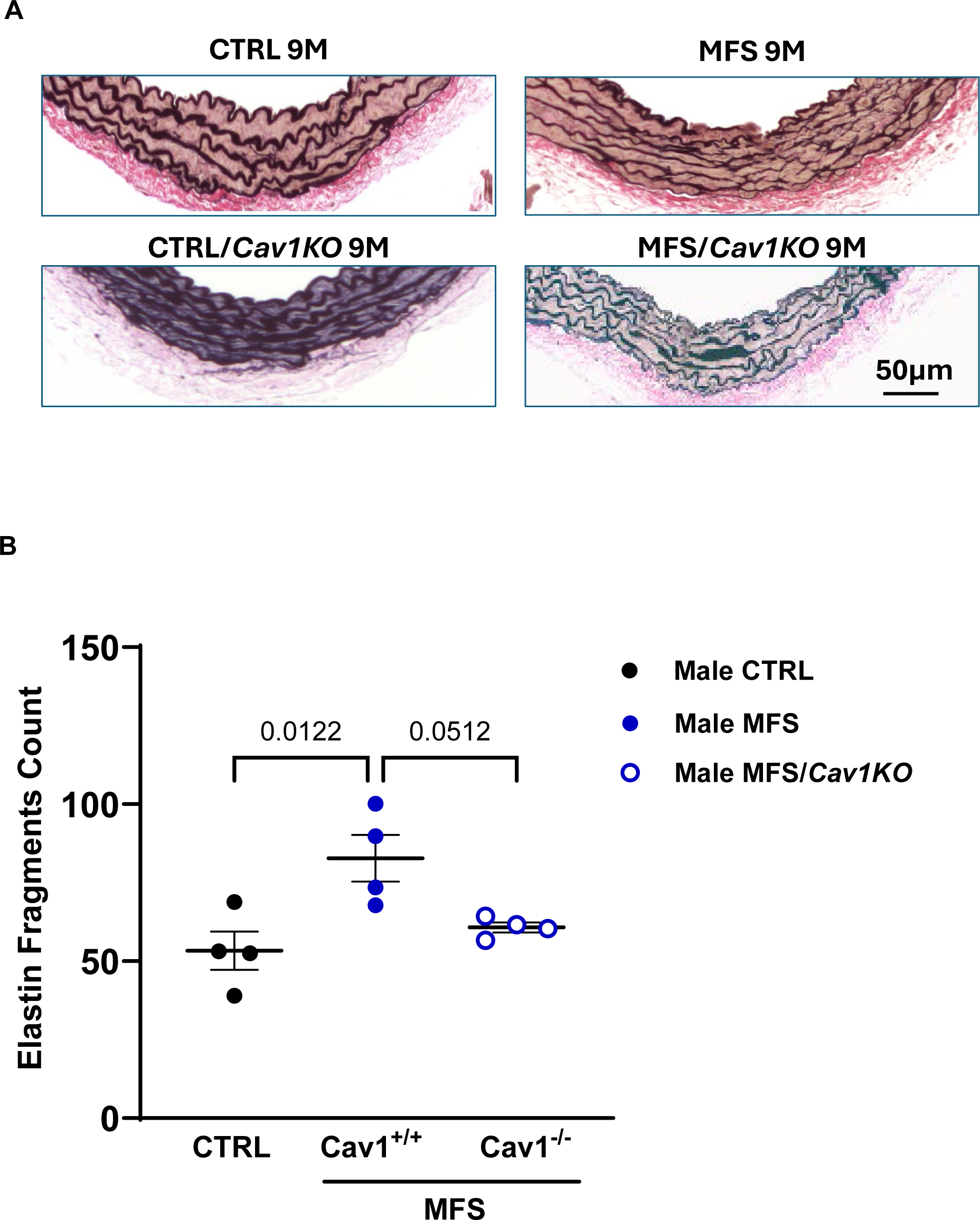
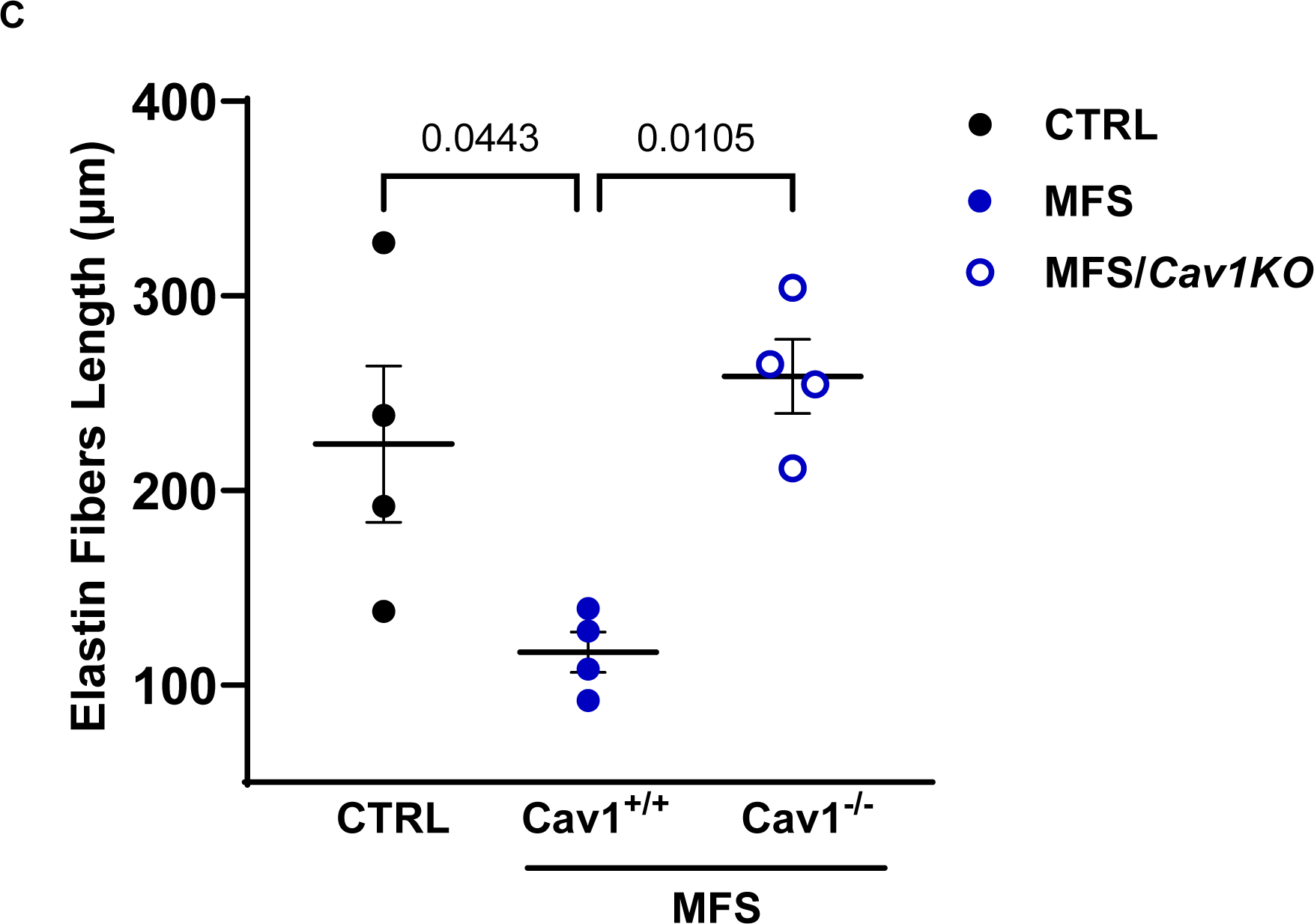
Aortic wall elastin structure is significantly improved in male MFS/*Cav1KO* mice. Data presents measurements of aortic wall elastin fragments counts and length within the aortic wall sections isolated from CTRL, MFS, and MFS/*Cav1KO* mice. **A)** Representative images showing Van Gieson staining of 10 µm aortic sections isolated from male CTRL, MFS, CTRL/*Cav1KO*, and MFS/Cav1KO mice, showing elastin in dark purple in the medial layer of mouse aortic wall (scale bar = 50µm). **B)** Quantification of elastin fiber counts shows a significant decrease in fragment counts in male MFS/*Cav1KO* aorta as compared to MFS mice. **C)** Similarly, *Cav1* deletion reduces elastin fiber fragments length in MFS mice, indicating a marked improvement in elastin structure within the aortic wall of MFS/*Cav1KO* mice. (Means ± SE, N = 4 mic/group, Two Way ANOVA followed by Tukey’s pairwise comparison, P ≤ 0.05).

## DISCUSSIONS

The present study aimed to investigate the potential role of Cav1 in regulating physiological responses pertinent to the progression of MFS-associated aortic root growth, aortic wall structural integrity, and aortic endothelial and smooth muscle function. Cav1 is a critical scaffolding protein predominantly found in the plasma membrane’s caveolae, which are small, flask-shaped invaginations rich in cholesterol and sphingolipids ^47,48^. Cav1 is integral to various cellular processes, including signal transduction, lipid regulation, inflammatory responses, and vesicular transportation ^29,49^, while directly interacting with a wide range of signaling proteins to modulate their activity ^50^. In addition to its role in normal cellular function, alterations in Cav1 expression and function are implicated in several pathophysiological conditions, such as inflammation ^51^, cancer ^52^, cardiovascular diseases ^53,54^, and diabetes ^55,56^. In the vasculature, Cav1 interacts with many membrane signaling molecules such as G-protein coupled receptors, insulin receptors, and cholesterol via its Cav scaffolding domain (CSD), and most importantly its critical role in regulating NO production within the vascular walls ^57–59^. Notably, Cav1 negatively regulates eNOS within the vascular walls, thereby influencing vascular tone and endothelial function ^23,60^.

In patients with MFS, the most common cause of mortality and morbidity is aortic root aneurysm, dissection, and rupture. MFS is caused by mutations in the *Fbn1* gene, leading to increased TGF-β signaling, reduced smooth muscle contractility, and impaired endothelial function, all of which contribute to the development of aortic aneurysms ^1,8^. Previous studies indicate that Cav1 plays a significant role in regulating both eNOS and TGF-β signaling within the vascular wall, as well as in the development of angiotensin II (AngII)-induced abdominal aortic aneurysm (AAA) ^61–64^. However, the role of Cav1 in MFS-associated aortic root aneurysm has yet to be investigated.

Our data clearly show that *Cav1* deletion induces a significant increase in endothelium-dependent relaxation in both male and female MFS mice, further confirming the inhibitory effects of Cav1 on endogenous endothelial NO production (**Fig. 3**). The EC_50_ values for acetylcholine in aortic rings show that the increase in vasorelaxation in MFS/*Cav1KO* aorta could be partially due to increased sensitivity of aortic endothelial cells to the vasorelaxant agent (**Fig. 3**). In line with this observation, we also found further marked decreases in PE-induced contraction in male MFS/*Cav1KO* mice aorta (**Fig. 5B**). Interestingly, in female MFS mice, Cav1 deletion did not impact PE-induced contraction (**Fig. 5B**). This may be explained by the observation that in female MFS mice, aortic contraction is already significantly lower compared to male MFS mice (**Fig. 5A**), reaching a threshold so low that is not further affected by the deletion of the *Cav1* gene. In both male and female CTRL and MFS aortic rings, pre-treatment with L-NAME increases aortic contraction. This phenomenon is expected as L-NAME blocks endogenous NO production within the aortic wall. The most interesting and important observation is that application of L-NAME increases MFS/*Cav1KO* mouse aortic contraction even further when compared to MFS mice (**Fig. 6C & 6D**). The effect of the general NOS inhibitor L-NAME on PE-induced contraction in both male and female MFS/*Cav1KO* mice further supports the notion that in the absence of the *Cav1* gene expression, endogenous NO levels are significantly elevated in aortic tissue, leading to a marked reduction in aortic contraction, and that this effect on aortic contraction can be reversed by L-NAME.

The interactions between Cav1 and nNOS have been explored in various studies. Cav1 can bind to nNOS, influencing its catalytic activity. This interaction is mediated by the Cav1 scaffolding domain, which can bind to the caveolin-binding motif present in nNOS. This binding typically results in the inhibition of nNOS activity, similar to the mechanism by which Cav1 interacts with eNOS ^49,65^. Dysregulation of the Cav1-nNOS interaction has been implicated in various diseases. For instance, in neurodegenerative disorders, the altered expression or function of Cav1 can lead to changes in nNOS activity, contributing to disease progression through increased oxidative stress and cellular damage ^66^. The interaction between Cav1 and eNOS is also a critical factor in regulating vascular function. In healthy blood vessels, Cav1 negatively regulates eNOS, reducing the production of NO, maintaining basal vascular tone and preventing excessive vasodilation ^49^. Under pathological conditions, the interplay between Cav1 and eNOS can be a key factor in disease progression. For example, in conditions such as hypertension or type 2 diabetes, elevated oxidative stress and inflammation, can lead to increases in *Cav1* expression, resulting in a significant decrease in NO bioavailability and production, therefore contributing to endothelial dysfunction, impaired myogenic tone, and increased vascular resistance ^56,67^.

In our study, the specific iNOS inhibitor 1400W had no impact on MFS/*Cav1KO* mouse aortic contraction (**Fig. 7**), leading to our conclusion that eNOS and/or nNOS are the most plausible sources for increased NO production within the aortic wall. Of these two potential candidates, several lines of evidence support the argument that eNOS is probably the primary target of *Cav1* deletion in our MFS study model. It has been consistently shown that eNOS is predominantly expressed in endothelial cells, playing crucial roles in normal endothelial function and vascular tone ^68^. In addition, the deletion of *Cav1* is known to disrupt its inhibitory interaction with eNOS, leading to increased eNOS activity and NO production ^22,51^.

On the other hand, nNOS is primarily expressed in nerve terminals, skeletal muscle, and to a lesser extent in vascular SMCs, and its contribution to systemic vascular NO production is typically less significant compared to eNOS ^69^. Our findings also align with previous studies that have demonstrated enhanced eNOS activity in the absence of Cav1, reinforcing the hypothesis that eNOS is the main source of the excessive NO production observed in MFS/*Cav1KO* mice ^70,71^. Further investigations using a more specific and targeted genetic manipulation of the *Cav1* gene would better elucidate details in the regulatory action of Cav1 on eNOS in the mouse model of MFS-associated aortic aneurysm.

Although reduction in aortic wall strength in MFS mice is evident in both sexes, the response to *Cav1* deletion seems to be sex specific in MFS/*Cav1KO* mice. In our study, we found that *Cav1* deletion enhances aortic wall strength only in male MFS, as evidenced by the improved aortic wall rupture points (**Fig. 8**). The observed difference could be due to different interactions between female and male sex hormones with pathways regulated by Cav1 protein. It is reported that Cav1 protein co-localizes and interacts with both the NH2-terminal and the ligand-binding domains of the androgen receptor (AR), and that downregulation of Cav1 can reduce AR activity ^72–74^. A strong body of evidence suggests that androgens can increase vascular SMC proliferation and hypertrophy, contributing to increased wall thickness ^75–77^. Androgens can also stimulate the synthesis of collagen in the vascular wall. Increased androgen levels have been associated with higher collagen deposition and reduced elastin content, leading to decrease elasticity ^78,79^. These reports may explain the observed protective effects of *Cav1* deletion on aortic wall strength and elastin structure in male MFS mice.

The sex-specific action of Cav1 deletion may also be explained by its interaction with estrogen receptor alpha (ERα). Cav1 can physically associate with estrogen receptor alpha (ERα) within caveolae leading to post-translational modifications that are necessary for the initiation of estrogen signaling pathways and its downstream beneficial effects on vascular function ^80,81^. In females, estrogen can modulate the expression and activity of various proteins involved in vascular function, including eNOS ^82^. It is possible that in female MFS mice, the presence of estrogen might mitigate the effects of *Cav1* deletion on aortic wall strength by differentially regulating pathways related to vascular remodeling and repair. Additionally, genes involved in ECM organization and inflammatory responses to injury (e.g. aortic root aneurysm) might be differently regulated in females, leading to different and sex-dependent outcomes following *Cav1* deletion. On the last note, although we have shown that *Cav1* deletion can increase aortic NO production in both male and female MFS mice, the downstream effects of enhanced NO signaling might differ in males and females. In males, the upregulated NO production may lead to more pronounced changes in ECM composition, thereby improving aortic wall strength, while in females, other regulatory mechanisms might dampen these effects. These proposed explanations should be further investigated and validated through carefully designed experiments in future studies.

Our findings also suggest that *Cav1* deletion could improve aortic wall elastin fiber structure within the aortic wall of MFS/*Cav1KO* mice (**Fig. 9**). The structural integrity of elastin fibers is crucial for supporting aortic wall elasticity. Elastin synthesis and structure can be directly or indirectly controlled by multiple signaling pathways originated from the ECM or cell-ECM interactions. In MFS, the *Fbn1* mutation can lead to an increase in TGF-β bioavailability and downstream signaling within the aortic wall. Excessive TGF-β signaling results in increased cell proliferation, differentiation, apoptosis, oxidative stress, and ECM degradation; all contributing to the development of multiple MFS manifestations ^5,83,84^. The TGF-β signaling pathway plays a significant role in the pathogenesis of MFS, in part due to the pathway’s role in regulating the expression of MMPs that control the normal turnover of the ECM components, playing important roles in connective tissue homeostasis, wound healing, and vascular wall integrity ^85–87^. In fact, in the experimental mouse model of MFS, MMP-2 and MMP-9 expression and activity are significantly elevated, causing increased elastolysis and elastin fragmentation within the aortic wall ^88–90^. The elastin deficiency and functional impairment has deleterious vascular effects including decreased vessel wall strength and elasticity that gives rise to high risk of mortality in MFS due to aortic dissection and rupture.

Interestingly, Cav1 is reported to also regulate the TGF-β signaling pathway, where the TGF-β type 1 receptor is endocytosed and degraded causing a decrease in downstream activation ^61,91^. Other groups have shown that crosstalk with the angiotensin II (AngII) type 1 receptor (AT1R) pathway can also weaken the vascular wall by increasing the bioavailability of TGF-β through a mitogen activated protein kinase ERK1/2-dependent signaling pathway, exacerbating the downstream detrimental effects ^92,93^. The inhibitory effect of Cav1 on both TGF-β and AT1R downstream signaling pathways can reduce the inflammation resulting from the secondary cascade and the downstream release of MMPs, which are the main players responsible for elastin fragmentation and loss of aortic wall elasticity ^88^. Hence, we expected that *Cav1* deletion would exacerbate elastin fragmentation in the aortic wall in MFS/*Cav1KO* mice. Surprisingly, we found that in male MFS/*Cav1KO* mice, *Cav1* deletion caused a significant increase in aortic wall strength (as evidenced by an increase in wall rupture point) and an improvement in aortic wall elasticity (as evidenced by increases in elastin fiber length and reduction in elastin fragments count within the aortic wall), with no beneficial effects observed in female subjects (**Fig. 8**). The observation that *Cav1* knockout decreases elastin fragmentation in the aortic wall of male mice can be possibly linked to a combination of reduced MMP activity, enhanced structural integrity via TGF-β signaling, hormonal influences, and reduced inflammation, collectively contributing to the preservation of elastin and the overall integrity of the aortic wall in male mice. However, there is no current evidence to support such a notion.

In addition, such potential explanation contradicts previous reports suggesting that *Cav1* knockout would decrease TGF-β and MMPs expression, therefore, reducing elastin degradation through altered MMP activity ^94,95^. The unexpected and sex-dependent phenomenon in male MFS/*Cav1KO* mice raises new questions about the regulatory impact of Cav1 on downstream signaling pathways involved in ECM turnover including TGF-β and MMPs. It is plausible that in MFS male mice, the absence of *Cav1* alters the regulation of TGF-β signaling, potentially leading to a decrease in MMP production. This altered signaling may stabilize the ECM and reduce elastin fragmentation. This hypothesis requires validation through further molecular and pharmacological studies.

Based on our data, we can only conclude that *Cav1* deletion provides beneficial effects on endothelial and SMC function in both male and female MFS mice, while improving aortic wall strength and elasticity only in male subjects (**Fig. 10**). It is also important to point out that despite the observed beneficial effects of *Cav1* deletion on endothelial and SMCs function, we did not see any beneficial and protective effects on aortic root growth, raising the questions about other regulatory mechanisms that contribute to the development of aortic root aneurysm in MFS mice.

**Figure 10.**
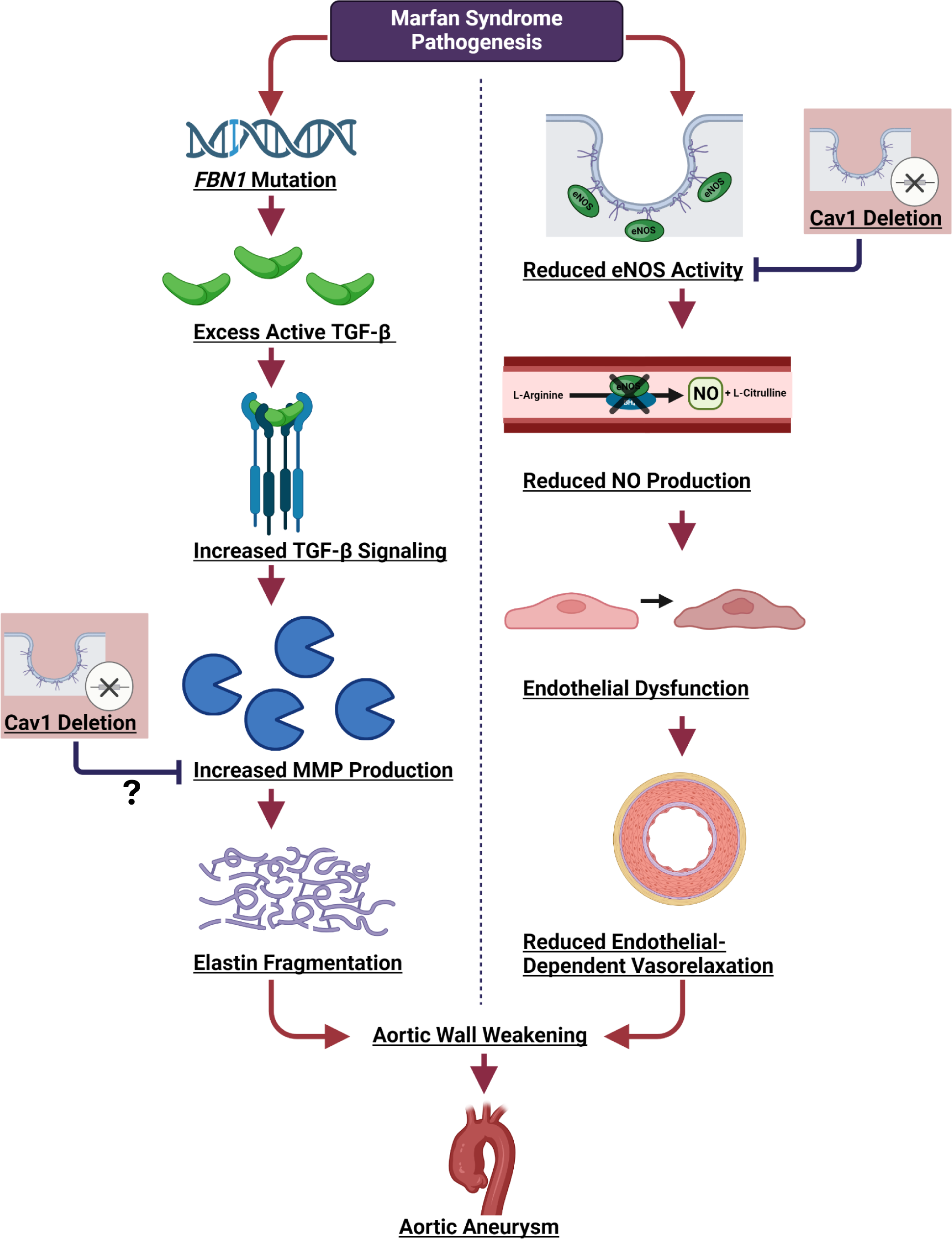
Potential impact of *Cav1* ablation on MFS aeropathy This schematic illustrates a proposed mechanism by which Marfan syndrome (MFS) could lead to aortic aneurysm, emphasizing the potential role of Caveolin-1 (Cav1). A mutation in the *FBN1* gene could result in an excess of active Transforming Growth Factor-beta (TGF-β). This increase in TGF-β signaling enhances the production of matrix metalloproteinases (MMPs), which degrade extracellular matrix components, causing elastin fragmentation and subsequent weakening of the aortic wall. Increased TGF and MMPS activity is known to lead to endothelial dysfunction, which is associated with a decrease in endothelial nitric oxide (eNOS) activity and endogenous nitric oxide (NO) production. Since Cav1 is known to have an inhibitory effect on eNOS activity, it is expected that the deletion of *Cav1* in MFS mice could lead to an improvement in endothelial function and NO production. This proposed mechanism underscores the importance of Cav1 in regulating key pathways involved in MFS pathogenesis and highlights the need for further investigation to validate these findings.

The current study has several limitations. The use of a mouse model and the specific experimental approach do not account for potential compensatory mechanisms by other isoforms of Caveolin, such as Cav2 and Cav3. Additionally, the systemic manipulation of *Cav1* gene expression in this study does not target a specific tissue of interest such as aortic wall endothelium or smooth muscle layer. A tissue-specific approach would provide a more detailed understanding of Cav1’s role in specific cell types and their contribution to MFS-associated aortic aneurysm. Future research should also incorporate more specific measurements of TGF-β and MMP signaling in the MFS/*Cav1KO* mice. This could involve detailed quantification of TGF-β levels and MMP activity in various tissues to better elucidate the mechanistic pathways by which Cav1 influences these signaling molecules in the context of MFS aneurysms. Such targeted studies will help clarify the role of Cav1 in regulating ECM dynamics and aeropathy in MFS. In addition, *in vivo* studies using ultrasound imaging should be conducted to determine the impact of *Cav1* deletion on the progression and severity of aortic aneurysms at different time points alongside cardiac function and structure. Ultrasound imaging can provide real-time, non-invasive assessments of aortic dimensions and structural integrity, offering valuable insights into the dynamic changes occurring in the aortic wall following *Cav1* deletion. Also, further studies assessing adapter proteins such as Cavin-1, which is essential for the formation and maintenance of caveolae and various cellular processes including signal transduction, endocytosis, and actin filaments and cytoskeletal interactions, may help explain the observed beneficial impact of *Cav1* knockout on aortic wall strength and elastin fragmentation. Finally, this study only investigates the effects of *Cav1* deletion on aortic root structure and function in 9-month-old CTRL and MFS mice. A longitudinal study that includes different age groups will further clarify the timeline for the observed protective effects.

In conclusion, this report is the first investigation of the potential regulatory role of Cav1 in the mouse model of MFS with a focus on the impact on endothelial and smooth muscle function (**Fig. 10**). The study of Cav1 in the context of MFS-associated aortic aneurysm is not only important for a deeper understanding of the disease’s pathophysiology but also holds promise for innovative treatments. Cav1’s role in modulating signaling pathways such as TGF-β and MMPs, both of which are critical in the pathogenesis of aortic aneurysms, highlights its potential as a therapeutic target. By bridging the gap between molecular insights and clinical applications, future research on Cav1 could pave the way for more effective management and improved outcomes for patients with MFS.

## ACKNOWLEDGEMENTS

We thank Mr. David Lowry at the Arizona State University Core Research Facility for his assistance with transmission electron microscopy imaging. This study was supported by a Faculty Grant [to M.E.] from the National Marfan Foundation.

## AUTHOR CONTRIBUTIONS

T.C. helped with designing and performing experiments, analyzing the data, and writing the manuscript. B.G. assisted with animal breeding, and data analysis. R.P. and T.B.J. helped with histological staining, image acquisition, image analysis, and manuscript editing. R.D. helped with myograph experiments and data collection. N.J. helped with experimental design, statistical analyses, and manuscript editing. J.V.E. helped with experimental design, data interpretation, manuscript editing, and aortic tissue preparation. M.E. was responsible for experimental design, data analysis, data presentation, manuscript writing and editing, project supervision, resources, and funding.

## CONFLICTS OF INTEREST

The author declares no conflicts of interest related to this study.

## DATA AVAILABILITY STATEMENT

The data presented in this study will be available on request from the corresponding author.

## Notes

### Competing Interest Statement

The authors have declared no competing interest.

