## Supplementary Figures for "Genetic Manipulation of Caveolin-1 in the Mouse Model of Marfan Syndrome Associated Aortic Root Aneurysm: Effects on Endothelial and Smooth Muscle Function"

**Supplementary Figure S1**

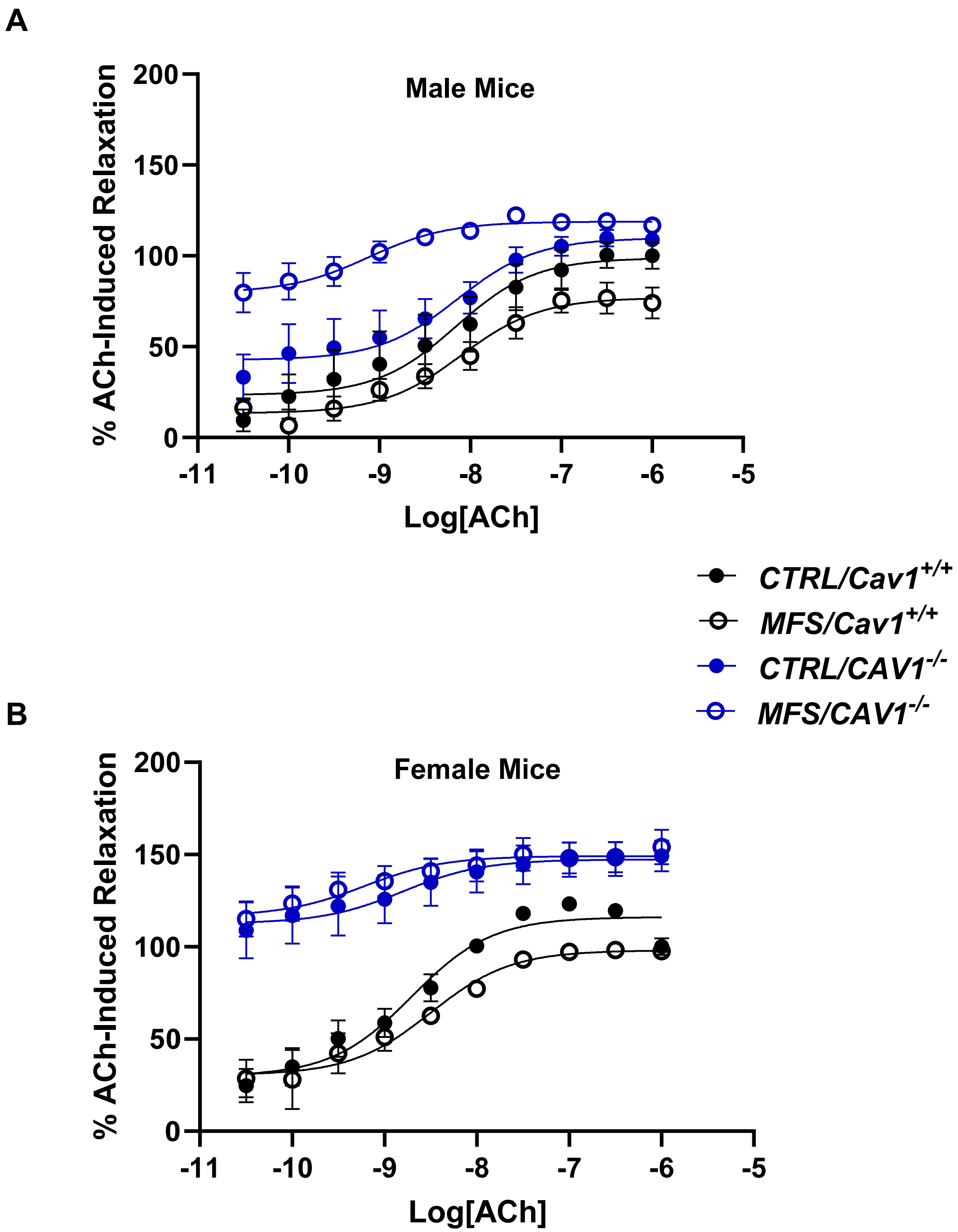


**Figure S1.** Dose-response curves (50 pM-1 µM) for acetylcholine (Ach)-induced aortic relaxation in aortic rings isolated from 9-month-old male **(A)** and female **(B)** CTRL, MFS, CTRL/Cav1KO, and MFS/Cav1KO mice. The $E_{max}$ and $EC_{50}$ values for Ach were derived from shown dose-response curves. (N = 9-12 mice/group).

**Supplementary Figure S2**

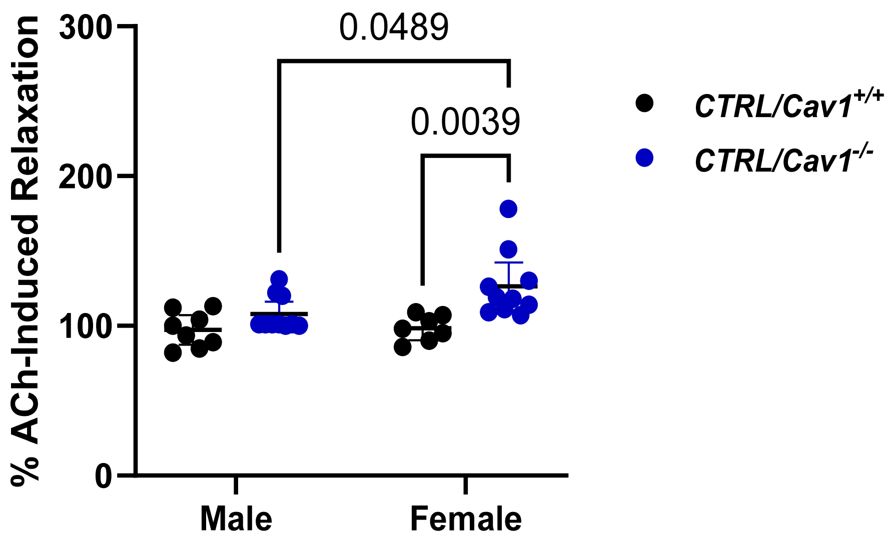


**Figure S2.** Maximum relaxation achieved in response to sub-max concentration of acetylcholine (500 nM) in aortic rings isolated from male and female CTRL and CTRL/*Cav1KO* mice. Data shows that *Cav1* deletion increased aortic relaxation only in female CTRL mice, indicating a sex-dependent effect. (Means ± SE, N = 8-11 mic/group, Two Way ANOVA followed by Tukey's pairwise comparison, P ≤ 0.05).

**Supplementary Figure S3**

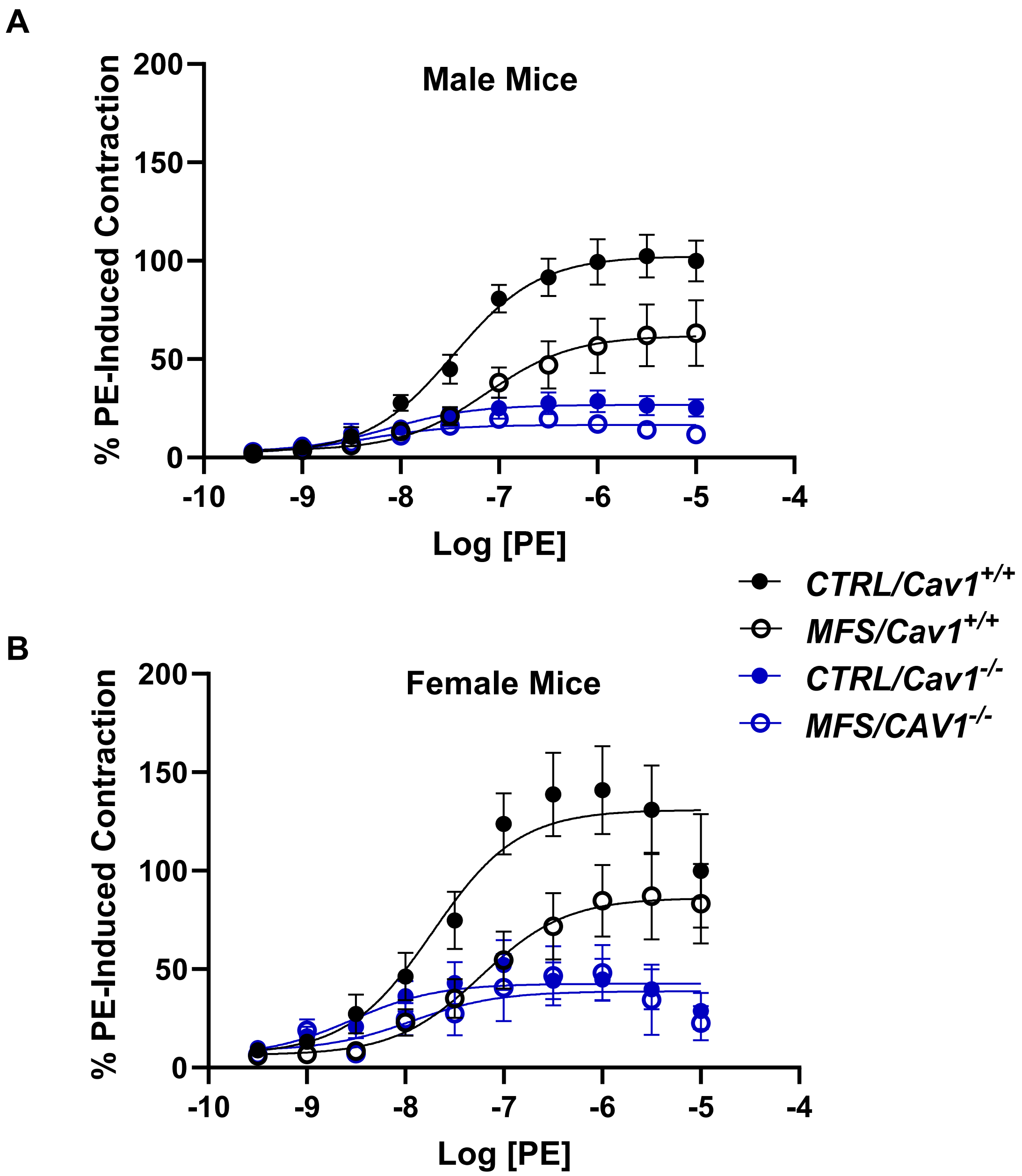


**Figure S3.** Dose-response curves (1 nM-50 μM) for phenylephrine (PE)-induced aortic contraction in aortic rings isolated from 9-month-old male **(A)** and female **(B)** CTRL, MFS, CTRL/Cav1KO, and MFS/Cav1KO mice. The values for $E_{max}$ and $EC_{50}$ were derived from shown dose-response curves. (N = 9-12 mice/group).

### Supplementary Figure S4

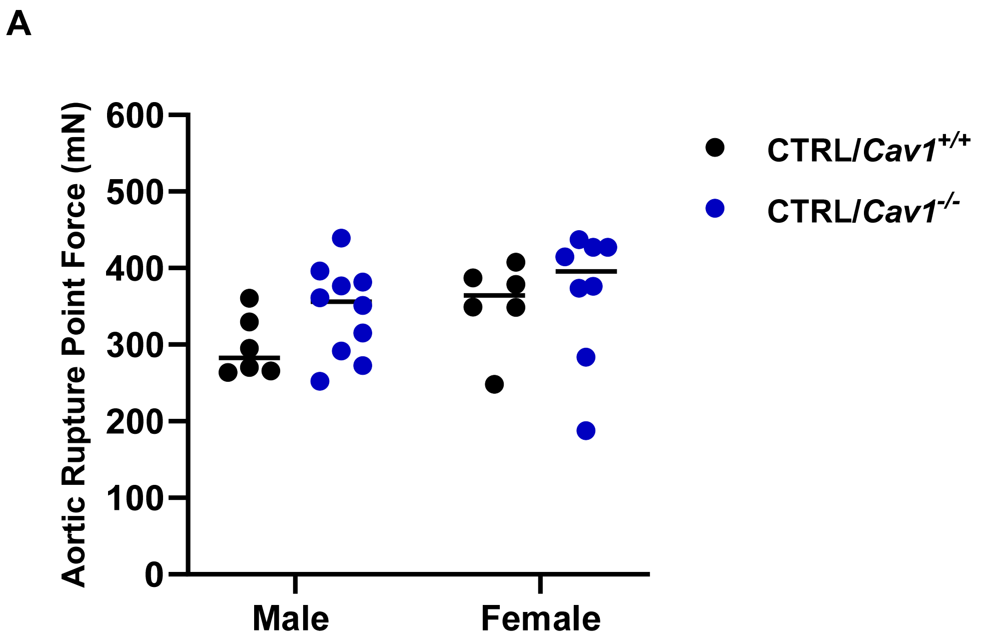


**Figure S4.** Maximum aortic wall strength (rupture point) in response to interval stretches of aortic rings in the myograph chamber. As shown, Cav1 deletion has no impact on aortic wall strength in male and female CTRL mice. (Means ± SE, N = 6-10 mic/group, Two Way ANOVA followed by Tukey’s pairwise comparison, $P \leq 0.05$).
